# Differential regulation of degradation and immune pathways underlies adaptation of the ectosymbiotic nematode *Laxus oneistus* to oxic-anoxic interfaces

**DOI:** 10.1101/2021.11.11.468236

**Authors:** Gabriela F. Paredes, Tobias Viehboeck, Stephanie Markert, Michaela A. Mausz, Yui Sato, Manuel Liebeke, Lena König, Silvia Bulgheresi

**Author notes:** Address correspondence to Silvia Bulgheresi,. Contributed equally to this work (Lena König, Silvia Bulgheresi).

## Abstract

Eukaryotes may experience oxygen deprivation under both physiological and pathological conditions. Because oxygen shortage leads to a reduction in cellular energy production, all eukaryotes studied so far conserve energy by suppressing their metabolism. However, the molecular physiology of animals that naturally and repeatedly experience anoxia is underexplored. One such animal is the marine nematode *Laxus oneistus*. It thrives, invariably coated by its sulfur-oxidizing symbiont *Candidatus* Thiosymbion oneisti, in anoxic sulfidic or hypoxic sand. Here, transcriptomics and proteomics showed that, whether in anoxia or not, *L. oneistus* mostly expressed genes involved in ubiquitination, energy generation, oxidative stress response, immune response, development, and translation. Importantly, ubiquitination genes were also upregulated when the nematode was subjected to anoxic sulfidic conditions, together with genes involved in autophagy, detoxification and ribosome biogenesis. We hypothesize that these degradation pathways were induced to recycle damaged cellular components (mitochondria) and misfolded proteins into nutrients. Remarkably, when *L. oneistus* was subjected to anoxic sulfidic conditions, lectin and mucin genes were also upregulated, potentially to promote the attachment of its thiotrophic symbiont. Furthermore, the nematode appeared to survive oxygen deprivation by using an alternative electron carrier (rhodoquinone) and acceptor (fumarate), to rewire the electron transfer chain. On the other hand, under hypoxia, genes involved in costly processes (e.g., amino acid biosynthesis, development, feeding, mating) were upregulated, together with the worm’s Toll-like innate immunity pathway and several immune effectors (e.g., Bacterial Permeability Increasing proteins, fungicides).

In conclusion, we hypothesize that, in anoxic sulfidic sand, *L. oneistus* upregulates degradation processes, rewires oxidative phosphorylation and by reinforces its coat of bacterial sulfur-oxidizers. In upper sand layers, instead, it appears to produce broad-range antimicrobials and to exploit oxygen for biosynthesis and development.

## INTRODUCTION

Fluctuations that lead to a decrease in oxygen availability are common in nature (Hermes-Lima and Zenteno-Savin, 2002). The physiological and behavioral response to oxygen deprivation has been studied in animals that naturally experience oxygen deprivation, such as frogs, goldfish, and turtles (Hochachka et al., 1996; 1997, 2001; Hermes-Lima and Zenteno-Savin, 2002), as well as in invertebrate genetic models (Clegg 1997; Nystul et al., 2003; Teodoro and O’Farrell, 2003; Haddad 2006). When oxygen deprived, these organisms must face the challenge of a drastic drop in ATP (the energy-storing metabolite adenosine triphosphate) production, which leads to the failure of energy-demanding processes that are crucial for maintaining cellular homeostasis. Anoxia-tolerant organisms, however, are capable to save energy by stopping energy-costly cellular functions (e.g., protein synthesis, ion pumping, cell cycle progression), maintain stable and low permeability of membranes, and produce ATP by anaerobic glycolysis (Hochachka et al., 1996; Teodoro and O’Farrell, 2003; Liu & Simon, 2004; Liu et al., 2006; Galli et al., 2014).

When parasitic and free-living nematodes, including the model organism *Caenorhabditis elegans*, are experimentally exposed to anoxia (<0.001 kPa O_2_), the intracellular ATP/ADP ratio drops dramatically and, within 10 h, they enter a state of reversible metabolic arrest called *suspended animation*. Namely, they stop to eat, move, develop or lay eggs, implying that oxygen deprivation affects their growth and behavior (Van Voorhies et al., 2000; Padilla et al., 2002; Nystul and Roth, 2004; Powell-Coffmann 2010; Fawcett et al., 2015; Kitazume et al., 2018). If these effects can be reversed upon oxygen reestablishment, the latter can also provoke a massive and sudden production of reactive oxygen species (ROS) that may overwhelm the organism’s antioxidant defense, and cause its death (reviewed in Hermes-Lima and Zenteno-Savin, 2002). Of note, an increase of mitochondrial ROS production was also observed in worms under hypoxia, because of the inefficient transfer of electrons to molecular oxygen (Nystul and Roth, 2004; Kim and Jin, 2015).

Because oxygen diffuses slowly through aqueous solutions, sharp concentration gradients of this electron acceptor may occur in marine environments and wet soil (Denny et al., 1993; Fawcett et al., 2015). It is at oxic-anoxic interfaces of marine sands that free-living nematodes coated with sulfur-oxidizing Gammaproteobacteria (Stilbonematinae) abound (Ott et al., 1989, 1991; Schiemer et al., 1990; Paredes et al., 2021). However, up to this study, the molecular mechanisms allowing symbiotic nematodes to withstand anoxia, and the inherent stress it is known to inflict upon metazoans, were unknown. Here, we incubated *Laxus oneistus* (Ott et al., 1995) in conditions resembling those it encounters in its natural environment (i.e. anoxic sulfidic or hypoxic), and applied comparative transcriptomics, proteomics and lipidomics, to understand how it copes with oxygen deprivation. Contrarily to our expectations, in anoxic sulfidic water *Laxus oneistus* did not appear to enter suspended animation. However, it upregulated genes required for ribosome biogenesis and energy generation, and for degradation pathways (e.g., ubiquitination-proteasome systems, autophagy) likely involved in recycling damaged cellular components and misfolded proteins into nutrients. Notably, under anoxic sulfidic conditions, it also upregulated putative symbiont- binding molecules such as lectins. In the presence of oxygen, on the other hand, the worm appeared to overexpress genes involved in energy-demanding processes (e.g., amino acid synthesis, development, feeding, and mating) and upregulated the synthesis of broad-range antimicrobials, likely via triggering the Toll/NF-kB pathway.

## RESULTS AND DISCUSSION

### The nematode *Laxus oneistus* did not enter suspended animation upon 24 h anoxia

To survive anoxia, nematodes enter suspended animation to suppress metabolism and conserve energy. The most notorious sign of suspended animation is the arrest of motility (Nystul et al., 2003; Chan et al., 2010; Kitazume et al., 2018).

Surprisingly, although the whole population of four tested nematode species, including *C. elegans*, was reported to be in suspended animation upon 10 h in anoxia (Kitazume et al., 2018), *L. oneistus* kept moving not only after 24-h-long incubations, but also upon 6-day-long incubations in anoxic seawater (three batches of 50 worms were incubated under each condition). Additionally, the symbiotic nematodes appeared morphologically normal (Supplemental movies 1-4).

The fact that we could not observed suspended animation, led us to hypothesize that *L. oneistus* evolved different strategies to survive oxygen deprivation.

### Stable transcriptional profile under hypoxic or anoxic sulfidic conditions

To understand the molecular mechanisms underlying *L. oneistus* response to oxygen, we subjected it to various oxygen concentrations. Namely, nematode batches were incubated under either normoxic (100% air saturation; O), hypoxic (30% air saturation; H) or anoxic (0% air saturation; A) conditions for 24 h. Additionally, given that *L. oneistus* thrives in reduced sand containing up to 25 µM sulfide (Ott and Novak., 1989; Paredes et al., 2021), we also incubated it in anoxic seawater supplemented with < 25 µM sulfide (anoxic sulfidic condition; AS).

While transcriptional differences of its symbiont (*Candidatus* Thiosymbion oneisti), incubated under normoxic (O) and hypoxic (H) conditions were negligible (Paredes et al., 2021), the expression profiles of nematode batches incubated under O conditions varied so much that they did not cluster (Figure S1). Consequently, there was no detectable differential expression between the transcriptomes of O nematodes and any of the other transcriptomes (H, A or AS; Figure S1B, C). We attribute the erratic transcriptional response of *L. oneistus* to normoxia to the fact that this concentration is not typically experienced by *L. oneistus* (Ott et al., 1989; Paredes et al., 2021).

As for the expression profiles of nematodes subjected to the H, A or AS conditions, replicates of each condition behaved more congruently (Figure S1B). While we did not find any significant difference between the A and AS nematodes, only 0.05% of the genes (8 genes; Data S1) were differentially expressed between the H and A nematodes and there was no significant difference between the H and A proteomes (t-test, FDR, Benjamini-Hochberg correction, p < 0.05; Figure S2A, Data S1). However, 4.8% of the expressed genes (787 out of 16,526) were differentially expressed between H and AS nematodes, with 434 upregulated under AS and 353 genes upregulated under H conditions (Figure S1C, Data S1).

Collectively, our data suggests that *L. oneistus* may be ill-equipped to handle normoxic sediment, but it maintains a largely stable physiological profile under both hypoxic and anoxic sulfidic conditions. Before discussing the subset of biological processes differentially upregulated in AS versus H nematodes and vice versa, we will present the physiological processes the worm appears to mostly engage with, irrespectively of the environmental conditions we experimentally subjected it to.

### Top-expressed transcripts under all tested conditions

To gain insights on *L. oneistus* basal physiology, we treated all the 16 transcriptomes as biological replicates (i.e., O, H, A and AS transcriptomes were pooled) and identified the 100 most abundant transcripts out of 16,526 based on functional categories extracted from the UniProt database (2021) and comprehensive literature search (Figure 1, Data S2). Our manual classification was supported by automatic eggNOG classification (Data S2). Similarly, the H and A proteomes were pooled, and the 100 most abundant proteins out of 2,626 were detected (Figure S2).

**Figure 1.**
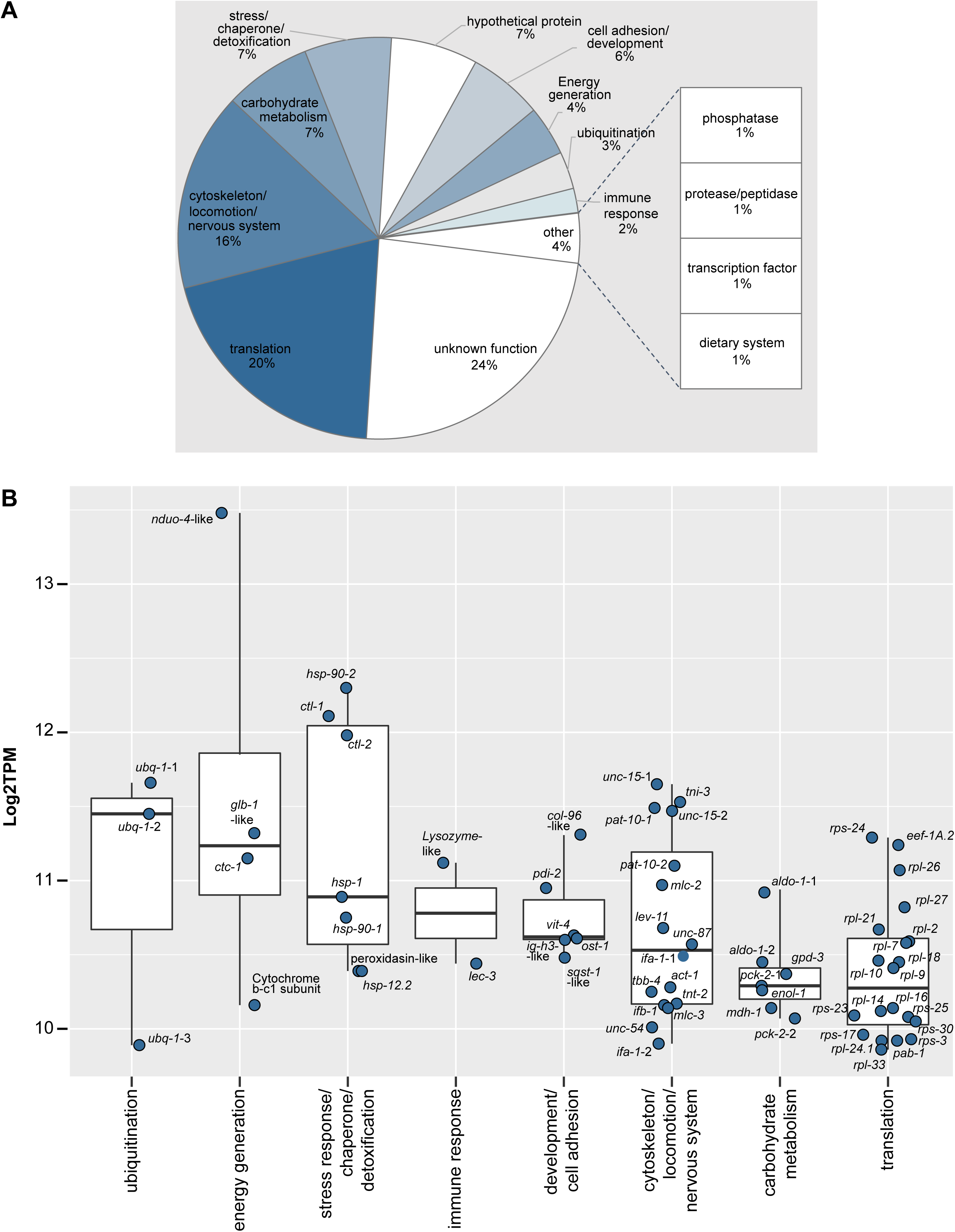
Relative transcript abundance and expression levels of the top 100 expressed genes of *L. oneistus* across all conditions. (A) Relative transcript abundance (%) of the top 100 expressed genes with a manually curated functional category. The top 100 expressed genes were collected by averaging the expression values (log_2_TPM) across all replicates of all incubations (Figure S1A, Data S1, and S2). Functional classifications were extracted from the curated database UniProt and from comprehensive literature search focused mainly on *C. elegans,* and confirmed with the automatic annotated eggNOG classification (Data S1). (B) Median gene expression levels of selected *L. oneistus* manually annotated functional categories of the top 100 expressed genes. Metabolic processes include both differentially and constitutively expressed genes. Each dot represents the average log2TPM value per gene across all replicates of all incubations. All gene names (or locus tags for unidentified gene names) are listed in Data S2.

Based on median gene expression values of the top 100 expressed genes, we found that some of the processes *L. oneistus* mostly engages with were ubiquitination (*ubq-1*, Stringham et al., 1992), energy generation (globin *glb-1*-like (Geuens et al., 2010), cytochrome c oxidase I subunit *ctc-1* (UniProtKB P24893), *nduo-4*-like (UniProtKB P24892), stress response and detoxification (e.g., *hsp-1*, *hsp-90*, *hsp12.2*, and catalases *ctl-1* and *ctl-2*; Birnby et al., 2000; Chávez et al., 2007), and immune defense (lysozyme-like proteins and *lec-3*) (Figure 1, Data S2).

Lastly, 48 out of the top 100 most expressed genes, were also detected among the top 100 proteins (Figure 1, Figure S2, and Data S2, Supplemental material). Despite the modest correlation between transcript and protein expression levels (r = 0.4) (Figure S3), there was an overlap in the detected biological processes (e.g., energy generation, stress response or detoxification categories, carbohydrate metabolism, cytoskeleton, locomotion, nervous system) (Figure S2).

All in all, except for those encoding for immune effectors, top-transcribed *L. oneistus* genes could not be ascribed to its symbiotic lifestyle. This differs to what observed for other chemosynthetic hosts, such as giant tubeworms and clams. Indeed, it is perhaps because these animals acquire their symbionts horizontally and feed on them as they are housed in their cells (and not on their surface) that they were found to abundantly express genes involved in symbiont acquisition, proliferation control and digestion (Sun et al., 2017, Hinzke et al., 2019; Yuen et al., 2019). Notably, we did observe a partial overlap of the most expressed gene categories (e.g., oxidative stress, energy generation, immune response), when *L. oneistus* was compared to the marine gutless annelid *Olavius* a*lgarvensis*. We ascribe the overlap to the fact that, albeit endosymbiotic, *O.* a*lgarvensis* also inhabits shallow water sand (Figure S4, Supplemental material) and, as hypothesized for *L. oneistus*, it may also acquire its symbionts vertically (Woyke et al., 2006; Dubilier et al., 2008; Wippler et al., 2016; Zimmermann et al., 2016).

To conclude, although both symbiont- (Paredes et al., 2021) and host-transcriptomics do not suggest a high degree of inter-partner metabolic dependence in the *L. oneistus* ectosymbiosis, the nematode seems well-adapted to both anoxic sulfidic (AS) and hypoxic (H) sand (Figure 2, Data S1). The transcriptional response of the worm to these two conditions is, however, significant (Figure 2, Data S1), and it will be reported next.

**Figure 2.**
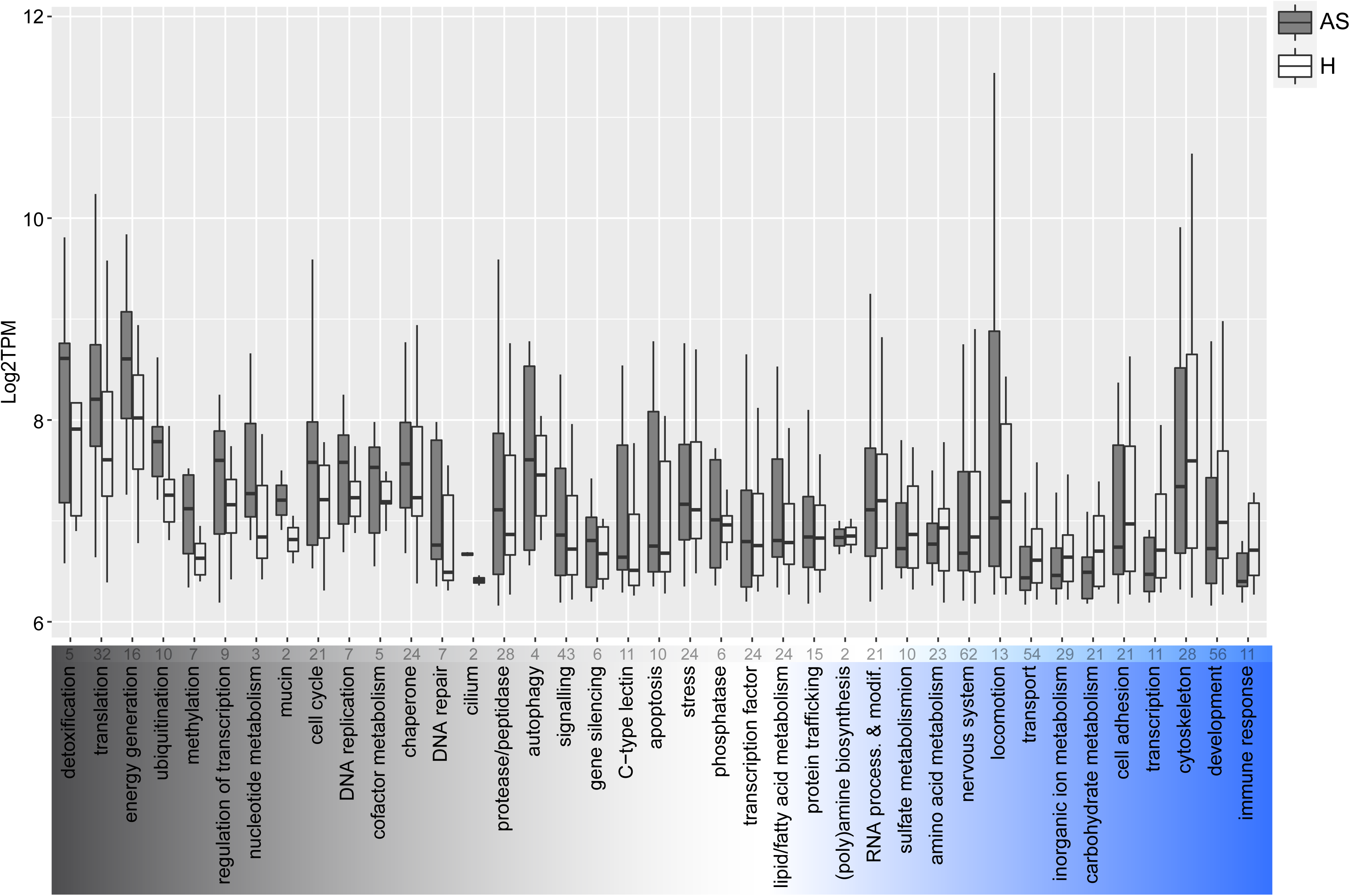
Median gene expression levels of selected *L. oneistus* metabolic processes among the differentially expressed genes between the hypoxic (H) and anoxic sulfidic (AS) conditions after 24 h. Individual processes among the differentially expressed genes are ordered according to their difference in median expression between the AS and H incubations. Namely, detoxification (far left) had the largest difference in median expression in the AS condition, whereas immune response (far right) had the largest median expression difference in the H condition. The absolute number of genes are indicated at the top of each process. Metabolic processes were manually assigned and confirmed with the automatic annotated eggNOG classification. For specific gene assignments see Data S1. Some genes are present in more than one functional category and processes comprising only one gene are not displayed in the figure but listed in Data S1.

### Genes upregulated in anoxic sulfidic (AS) nematodes

#### Chaperones and detoxification

The expression of chaperone-encoding (e.g., *hsp12.2*, *grpE*, *dnaJ/dnj-2*, *pfd-1*, *pfd-6*; Naylor et al., 1996; Lundin et al., 2008; Bar-Lavan et al., 2016), and ROS-detoxifying-related genes (e.g., superoxide dismutase *sod-2* and a putative glutathione peroxidase, involved in the detoxification of superoxide dismutase and hydrogen peroxide, respectively; Suzuki et al., 1996; Margis et al., 2008) were higher in AS nematodes (Figures 2 and 3). Notably, transcripts encoding for the heme-binding cytochrome P450 *cyp*-13B1 were also more abundant in AS (Figure 3), perhaps to increase the worm’s capacity to cope with putative ROS formation (Oliveira et al., 2009). Indeed, as cells start being oxygen-depleted, mitochondrial ROS accumulate because of the inefficient transfer of electrons to molecular oxygen (Semenza, 1999; Nystul and Roth, 2004; Selivanov et al., 2009; Kim and Jin, 2015). Alternatively, the upregulation of antioxidant-related genes in AS worms could represent an anticipation response to an imminent reoxygenation. In animals alternating between anoxic and oxygenated habitats, the re-exposure to oxygen can be very dangerous, as it creates a sudden ROS overproduction that may overwhelm the organism’s oxidative defense mechanisms (Hermes-Lima and Zenteno-Savin, 2002; Hashimoto et al., 2004). Although it has not been reported for nematodes, overexpression of ROS-counteracting genes is consistent with what has been reported for vertebrates and marine gastropods which, just like *L. oneistus*, alternate between oxygen-depletion and reoxygenation (Hermes-Lima and Zenteno-Savin, 2002).

**Figure 3.**
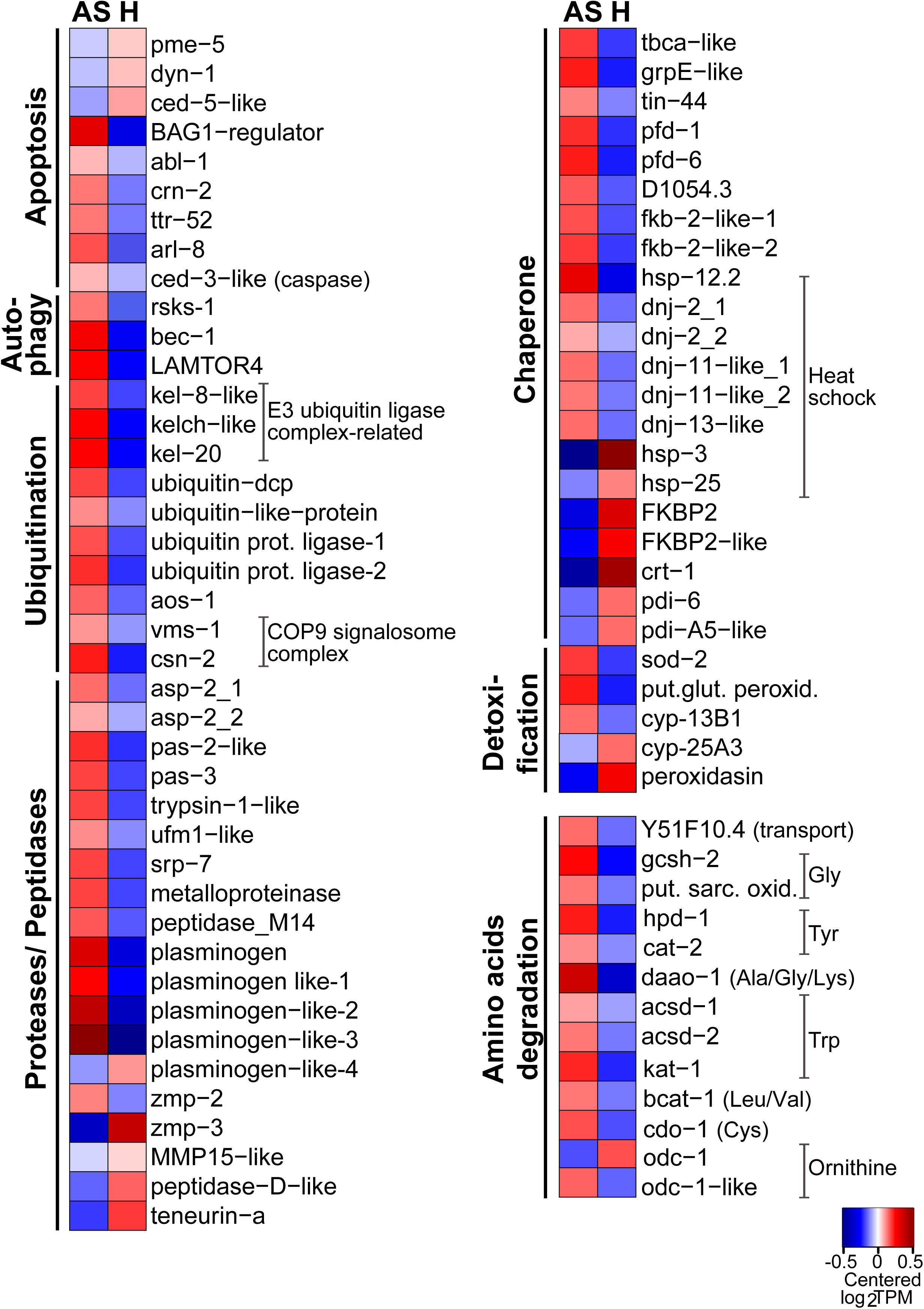
Genes involved in detoxification, ubiquitin-proteasome, autophagy, apoptosis, and amino acids degradation were predominantly expressed in AS worms. Heatmap displaying genes upregulated in AS (anoxic sulfidic) relative to H (hypoxic) worms after 24 h- long incubations under one of the two conditions (1.5-fold change, FDR ≤ 0.05). Expression levels are displayed as mean-centered log_2_TPM value (transcripts per kilobase million). Genes are ordered by function in their respective metabolic pathways. For each process, the minority of genes that were upregulated in H worms is shown in Data S1. Red denotes upregulation, and blue downregulation. Prot. protein, COP9: Constitutive photomorphogenesis 9. Dcp: domain-containing proteins. Put. glut. peroxid.: putative glutamate peroxidase. Put. sarc. oxid.: putative sarcosine oxidase.

#### Mitochondrial and cytoplasmic ribosome biogenesis

In the cellular stress imposed by oxygen deprivation, mitochondria are central to both death and survival (Borutaite et al., 1995; Brookes et al., 2004; Brenner et al., 2012; Hawrysh et al., 2013; Galli et al., 2014). In this scenario, calcium regulation, the scavenging of ROS or the suppression of their production, and/or inhibition of the mitochondrial permeability transition pore (MPTP) opening, might help to preserve mitochondrial function and integrity (Horwitz et al., 1994; Murphy et al., 2008; Galli et al., 2014; Fanter et al., 2020). In addition, removal of specific mitochondrial components (mitochondrial-associated protein degradation, MAD), might also arise to maintain the overall mitochondrial homeostasis (Chatenay-Lapointe and Shadel, 2010; Heo et al., 2010). Perhaps as a response to anoxia-induced stress (reviewed in Galli et al., 2014), a gene involved in MAD (*vms-*1) (Chatenay-Lapointe and Shadel, 2010; Heo et al., 2010), was upregulated in AS worms (Figure 4). More abundant in this condition were also transcripts encoding for mitochondrial transmembrane transporters *tin-44*, *slc-25A26* and *C16C10.1* (UniProtKB O02161, Q18934, Q09461), putatively transporting, peptide-containing proteins from the inner membrane into the mitochondrial matrix, such as S-Adenosyl Methionine (Figure 6). Surprisingly, although the translation elongation factor *eef-1A.2* (Tullet, 2015) was downregulated in AS worms, not only various mitochondrial ribosome structural components (28S: *mrps*, 39S: *mrpl*; Kaushal et al., 2014), and mitochondrial translation-related genes (e.g., *C24D10.6* and *W03F8.3*; Sharika et al., 2018) were upregulated in AS nematodes, but also several cytoplasmic ribosome biogenesis (40S: *rps*, 60S: *rpl;* Melnikov et al., 2012) and subunit assembly genes (e.g., RRP7A−like, You et al., 2015) (Figure 4).

**Figure 4.**
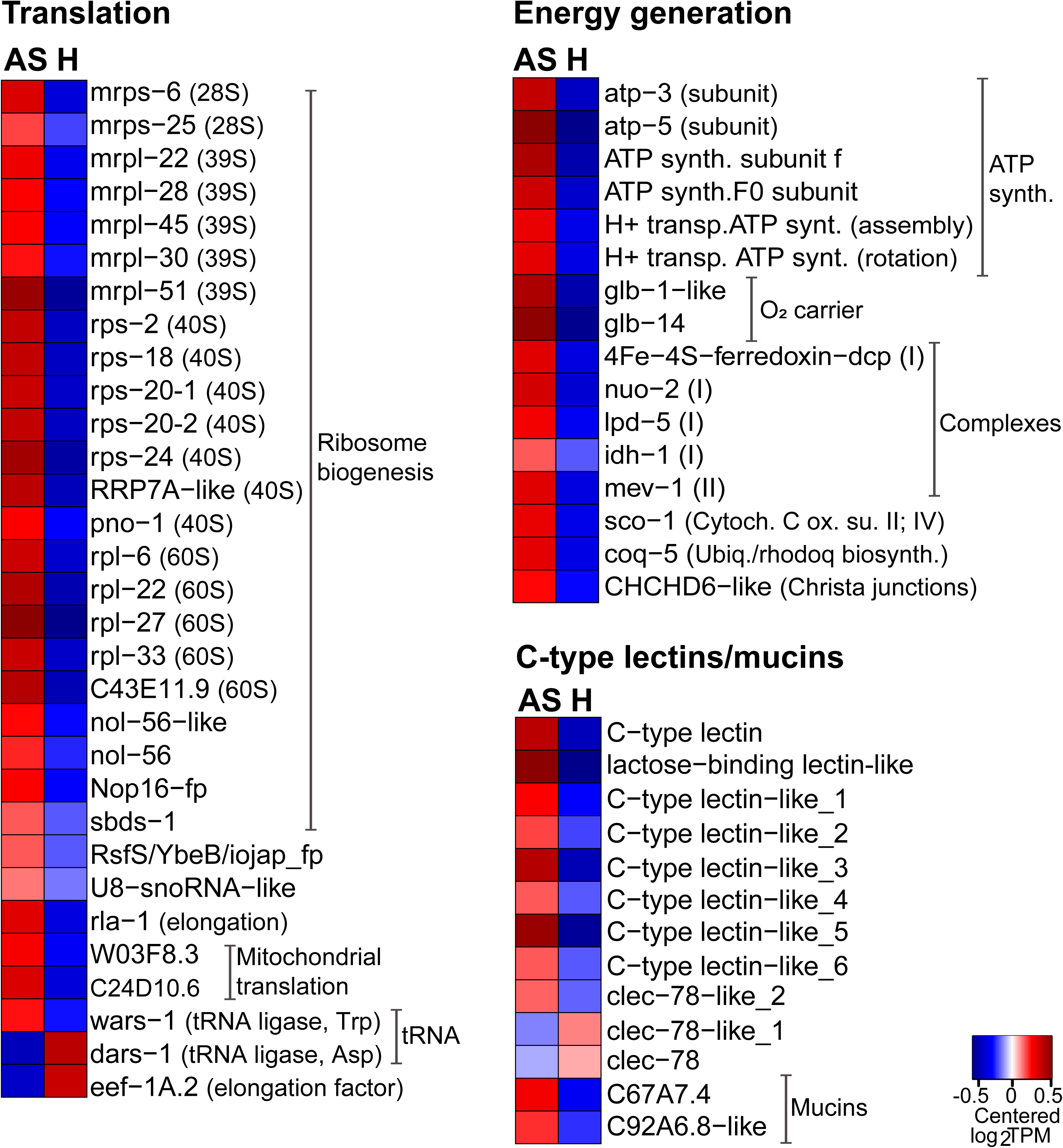
Genes involved in translation and energy generation and genes encoding for C-type lectins and mucins were predominantly expressed in AS worms. Heatmap displaying genes upregulated in AS (anoxic sulfidic) relative to H (hypoxic) worms, upon 24 h- long incubations under one of the two conditions (1.5-fold change, FDR ≤ 0.05). Expression levels are displayed as mean-centered log_2_TPM values (transcripts per kilobase million). Genes are ordered by function in their respective metabolic pathways. For each process, the minority of genes that were upregulated in H worms is shown in Data S1. Red denotes upregulation, and blue downregulation. Fp: family-containing protein. Cytoch. C ox. su. II.: cytochrome c oxidase subunit II. Ubiq./rhodoq biosynth.: Ubiquinone or rhodoquinone biosynthesis.

Taken together, the maintenance of mitochondrial homeostasis, an anticipatory response to a potential upcoming ROS insult (see Chaperones and detoxification section) and/or their involvement in extra-ribosomal functions (Chen et al., 2010; Savada et al., 2014; Xu et al., 2016) might explain the upregulation of ribosomal biogenesis-related genes in AS nematodes. Although upregulation of ribosomal proteins has also been observed in anoxic gastropods (Larade et al., 2001), increased ribosomal biogenesis (which oftentimes directly correlates with an increase of protein synthesis) is not expected in animals that must repress their metabolism to cope with oxygen deprivation (Thomas et al., 2000; Hochachka and Lutz 2001; Shukla et al., 2012).

#### Energy generation

Equally surprising was the upregulation of all differentially expressed genes related to energy generation in AS nematodes (Figure 4). Namely, besides putative oxygen-binding globulin-like genes (e.g., *glb-1*, *glb-14*, Geuens et al., 2010), the following were upregulated in AS nematodes: key structural genes (e.g., *atp-3*, *atp-5,* Xu et al., 2018), assembly-related genes (H+-transport ATP synthase, Maglioni et al., 2016) of the mitochondrial ATP synthase (complex V), genes related to complex I (*lpd-5*, *nuo-2,* McKay et al., 2003; Rea et al., 2007), a subunit of the succinate dehydrogenase involved in complex II (*mev-1,* Hartman et al., 2001), a mitochondrial cytochrome C oxidase subunit II assembly gene related to complex IV (*sco-1,* Williams et al., 2005), and a mitochondrial gene (*coq-5*), involved in the synthesis of either ubiquinone (Q, aerobic) or rhodoquinone (RQ, anaerobic) electron carriers (Buceta et al., 2019) (Figure 4). This suggests that, under anoxia, the electron transfer chain (ETC) is rewired in such way that electrons still enter the ETC at complex I, but instead of reaching complex III and IV they are transferred to RQ. This, in turn, shuttles the electrons to succinate dehydrogenase. The latter enzyme uses fumarate as an alternative electron acceptor, reducing it to succinate. This mechanism would maintain the flow of electrons through the ETC, and, it would prevent mitochondrial ATP generation (complex V) from shutting down (Buceta et al., 2019; Del Borrello et al., 2019).

In short, under AS, similarly to what has been observed in other free-living and parasitic nematodes, complex I appears to be the sole proton pump in this truncated form of ETC (Buceta et al., 2019; Del Borrello et al., 2019). In accordance with this hypothesis, tryptophan (Trp) degradation-related genes (*acsd-1*, *acsd-2*) and the Trp RNA ligase (*wars-1*; Tsai et al., 2017) that might be required to synthesize RQ (Buceta et al., 2019; Del Borrello et al., 2017; Tan et al., 2020) were upregulated under AS. Intriguingly, upregulated was also an isocitrate dehydrogenase gene (*idh-1*). This produces reducing equivalent (NADPH) carrying electrons that may fuel complex I (Smolková et al., 2012; Martínez-Reyes et al., 2020), but it might also add to the stimulation of the antioxidant capacity or to the maintenance of redox homeostasis by regenerating reduced glutathione (Hermes-Lima and Zenteno-Savin, 2002; Penkov et al., 2015; Yang et al., 2019).

If glycolysis is a key process for ATP generation in anoxia (Lutz et al., 1997; Semenza et al., 2001; Hochachka et al., 2001; Huang et al., 2008; Larade et al., 2009) and if, consistently, *hxk-2* was upregulated under this condition (Figure 6), based on the expression levels of transcripts encoding for alpha-amylases (see Carbohydrate metabolism in Figure 6), starch and/or glycogen (Jackson and McLaughlin, 2009) may be the prominent carbon sources under anoxic sulfidic conditions.

#### Ubiquitin-proteasome system and proteases

Proteolysis supplies amino acids or polypeptides to the cells, while impeding the accumulation of damaged or misfolded proteins. The two main mechanisms of cellular proteolysis are the lysosome-mediated intracellular protein degradation (autophagy) and the proteasome-mediated protein degradation (ubiquitin-proteasome system, UPS). In the latter, ubiquitin-protein ligases covalently attach ubiquitin to proteins, allowing their recognition and further degradation by the proteasome (Lodish et al., 2008; Papaevgeniou and Chondrogianni, 2014).

As shown in Figure 1, transcripts encoding for polyubiquitin (*ubq-1*), had the highest median gene expression across all transcriptomes. However, all ubiquitination-related genes detected in the differential gene expression analysis between the AS and H conditions, were upregulated in AS worms (Figure 2 and 3, Data S1). For example, *aos-1,* encoding for a subunit of the ubiquitin-activating enzyme (E1) (Jones et al., 2001), two ubiquitin-protein ligases (E3s without detected cullin domains; Papaevgeniou and Chondrogianni, 2014), and kelch-like genes (e.g., *kel-8*-like and *kel-20*). The former are BTB-domain containing proteins known to interact with E3 enzymes, with *kel-8* being involved in the degradation of glutamate neuroreceptors (Schaefer and Rongo 2006; Stogios et al., 2005; Kim et al., 2018). Additional ubiquitination-related genes upregulated in AS were *csn-2*, encoding for a component of the COP9 signalosome complex (Pintard et al., 2003; Brockway et al., 2014), and proteasome genes (*pas-2* and *pas-3;* Fraser et al., 2000; Blumenthal et al., 2002).

Among the proteases that were upregulated in AS worms, aspartyl proteases have been involved in neurodegeneration (Syntichaki et al., 2002), whereas plasminogen and the zinc matrix metalloproteinase ZMP-2 were both reported to mediate degradation of extracellular matrix (ECM) (Vassalli et al., 1991; Altincicek et al., 2010; Fischer, et al., 2014) (Figure 3). *C. elegans* ZMP-2 was also shown to prevent the accumulation of oxidized lipoproteins (Fischer et al., 2014), and, therefore it may contribute to the enhanced antioxidant response observed in this condition.

#### Autophagy and amino acid degradation

Besides acting coordinately to withstand stress, autophagy cooperates with apoptotic UPS for the recovery and supply of nutrients when these are scarce (Vabulas et al., 2005; Scott et al., 2004; Huber and Teis, 2016; reviewed in Wang RC et al., 2010 and Russel et al., 2014). Transcripts of two autophagy-related genes, *bec-1* (Liang et al., 1999) and the Ragulator complex protein LAMTOR4 (C7orf59-like) (Bar-Peled et al., 2012) were more abundant in AS nematodes (Figure 3). While the former positively regulates autophagy (Liang et al., 1999; Meléndez et al., 2003), the latter interacts with the mTOR Complex I (mTORC1), and tethers small GTPases (Rags and Rheb) to the lysosomal surface (Bar-Peled et al., 2012). When amino acid levels are low, mTORC1 is not translocated to the lysosomal surface (Wang et al., 2009; Bar-Peled et al., 2012), thereby favoring catabolic processes such as autophagy (Thompson et al., 2005). We propose that amino acid scarcity might result from the upregulation of genes involved in the degradation of lysin, glycin, tyrosin, cystein, leucin, isoleucin, valin or tryptophan (Figure 3, Data S1). This would decrease mTORC1 activity and, in turn, stimulates nutrient recycling via autophagy in AS worms.

Conversely, we hypothesize that in H worms, active mTORC1 interacts with the ribosomal protein S6 kinase (S6K), encoded by the *rsks-1* gene which is also up in H worms (Ladevaia et al., 2014) (Figure 3). This direct interaction, upon a cascade of phosphorylation events, would stimulate translation, and ultimately cell growth and proliferation (Ma et al., 2009, Howell et al., 2011, and Ladevaia et al., 2014).

All in all, although it is currently unclear whether increased autophagy is beneficial or detrimental, under AS conditions, the upregulation of genes involved in self-digestion might play a protective role and foster recovery from starvation (Thompson et al., 2005), pathogens (Huber and Teis, 2016) or from neuronal and muscular degeneration induced by oxygen deprivation (Murphy and Steenbergen 2008).

#### Lectins and mucins

Given that symbiont attachment may be mediated by Ca^2+^-dependent lectins (Nussbaumer et al. 2004, Bulgheresi et al., 2006, 2011) and given that, under anoxia, the symbiont appeared to proliferate more (Paredes et al., 2021), we expected nematode lectins to be upregulated under this condition. Indeed, nine C-type lectin domain (CTLD)-containing proteins were upregulated in AS *L. oneistus* adults and only two (*clec-78* and *clec-78*-like-2) were upregulated in the presence of oxygen (Figure 4). In addition to CTLD-containing proteins, mucins, a class of glycoproteins with more than 50% of its mass attributable to O-glycans, were also upregulated in AS nematodes. Considering that mucin glycans are used by vertebrate gut commensals for attachment, as well as a source of nutrients (Koropatkin et al., 2012), it is conceivable that their upregulation in anoxia (Figure 4), together with that of CTLD-containing proteins, would foster symbiont attachment.

We hypothesize that overexpression of two classes of putative symbiont-binding molecules, lectins and mucins, under conditions favoring symbiont proliferation (i.e., AS condition, Paredes et al., 2021) may mediate bacterial coat reinforcement.

#### Apoptosis

Mitochondria play an important role in apoptosis induction (Simon et al., 2000; Martínez-Reyes et al., 2020). Indeed, MPTP opening due to ROS (or the severe ATP decline imposed by the absence of oxygen) may cause cytochrome C release from mitochondria and this, in turn, triggers caspase activation (Martinou et al., 2000; Simon et al., 2000; Gogvadze et al., 2006; Galli et al., 2014). We observed that transcripts encoding for *sco- 1*, a gene needed for the synthesis and assembly of mitochondrial cytochrome C (Williams et al., 2005) were more abundant in AS worms (Figure 4). Further, we observed upregulation of Caspase-3 (*ced-3*) which belongs to a family of cysteine proteases involved in apoptosis (Mangahas et al., 2005; Kaufmann et al., 2008) and which is activated upon mitochondrial cytochrome C release into the cytosol (Liu et al., 1996; Tafani et al., 2000; Kaufmann et al., 2008; Martínez-Reyes et al., 2020). Additional apoptosis-related genes that appeared to be upregulated in AS worms were: *bec-1* (Figure 3), a gene that promotes autophagy and fine-tunes the Ced-3-mediated apoptosis (Liang et al., 1999; Takacs-Vellai et al., 2005); *ttr-52*, which mediates apoptotic cell recognition prior to engulfment (Wang, X. et al., 2010; Chen et al., 2013); a BAG family molecular chaperone regulator 1 (BAG1-regulator); a cell-death-related nuclease *crn−2* (Parrish et al., 2003; Samejima et al., 2005) and phagolysosome forming *arl-8* (Sasaki et al., 2013), and a tyrosine kinase Abl-1, (*abl-1*) that modulates apoptotic engulfment pathways (Hurwitz et al., 2009).

#### Lipid catabolism

Genes involved in lipid metabolism were similarly expressed between the AS and H conditions (Figure 2, Data S1). In accordance, lipidomes of nematodes incubated in the presence or absence of oxygen were not significantly different (Figure S5, Supplemental material). However, in line with the overall upregulation of degradation pathways, we observed upregulation of genes involved in FA beta-oxidation (*kat-1*; Berdichevsky et al., 2010), in lipid digestion (the lipase *lipl-6*; UniProtKB E2S7J2), and lipid degradation (a peripilin-2-like protein; Chughtai et al., 2015). Moreover, a gene that might be involved in oxidative-stress tolerance (a stearic acid desaturase *fat-7* regulating the first step of the fatty acid desaturation pathway (Horikawa et al., 2009) was also upregulated in AS worms. Lipid degradation under anoxia might be a strategy to overcome starvation (Krivoruchko and Storey, 2015).

Notably, we also observed an upregulation of two genes involved in phosphatidylcholine (PC) synthesis (*pmt-1*, *pmt-2*, Brendza et al., 2007) (Figure 5). Intriguingly, PC was more abundant in the anoxic symbiont (Paredes et al., 2021), although the latter cannot synthetize it. Thus, their upregulation in AS worms suggests worm-symbiont lipid transfer.

**Figure 5.**
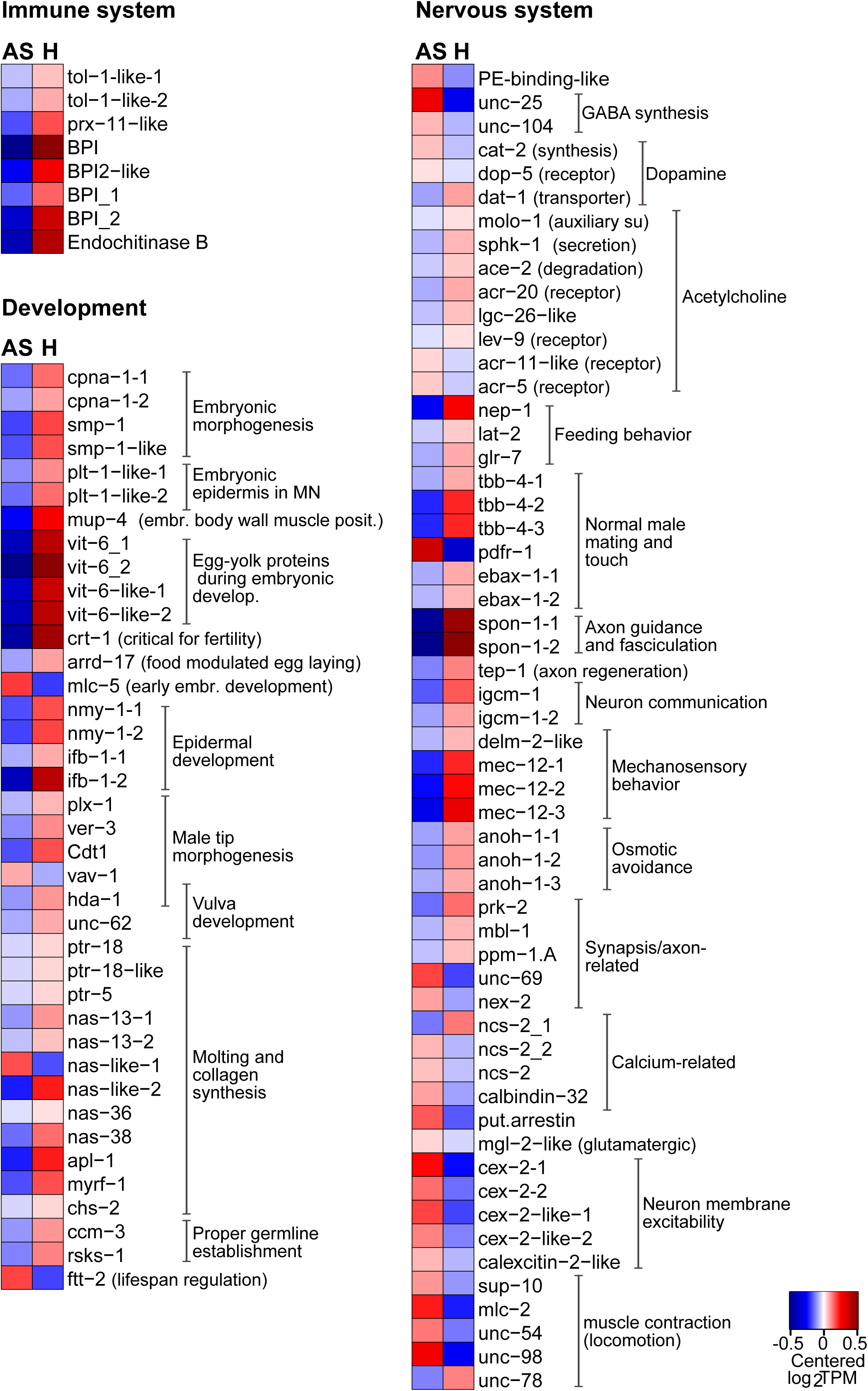
Genes involved in immune response, development and nervous system were predominantly expressed in hypoxic (H) worms. Heatmap displaying genes upregulated in H relative to AS worms, upon 24 h-long incubations under one of the two conditions (1.5-fold change, FDR ≤ 0.05). Expression levels are displayed as mean-centered log_2_TPM value (transcripts per kilobase million). Genes are ordered by function in their respective metabolic pathways. For each process, the minority of genes that were upregulated in AS worms is shown in Data S1. Red denotes upregulation and blue downregulation. MN: mechanosensory neurons. Embr. body wall muscle posit.: Embryonic body wall muscle positioning. Put.: putative.

#### GABA- and glutamate-mediated neurotransmission

Upregulated genes related to GABA synthesis were, *unc-25*, *unc-104* and *pdxk-1* (pyridoxal phosphate hexokinase) (Thomas et al., 1990; Mclntire et al., 1993; Jin et al., 1999; Gally et al., 2003; Nordquist et al., 2018; Risley et al., 2016) (Figure 5, Data S1). Consistent with an expected increase in glutamate requirement as a direct GABA precursor (Martin et al., 1993), we observed downregulation of two glutamine synthetases and a delta-1-pyrroline-5-carboxylate synthase (*gln-3* and *alh-13* respectively; van der Vos et al., 2012; Yen et al., 2021; Figure 6), known to convert glutamate to glutamine or to proline, respectively. Furthermore, an *mgl-2* like gene encoding for a glutamate receptor, which is activated in the presence of glutamate (Tharmalingam et al., 2012), was up in AS worms. Note that, when oxygen is limited, glutamate may act as a neurotoxic amino acid (Baker et al., 1991; Lutz et al., 2003a). Therefore, increased GABA biosynthesis might, beneficially, prevent its accumulation (Milton et al., 2002; Mathews et al., 2003).

**Figure 6.**
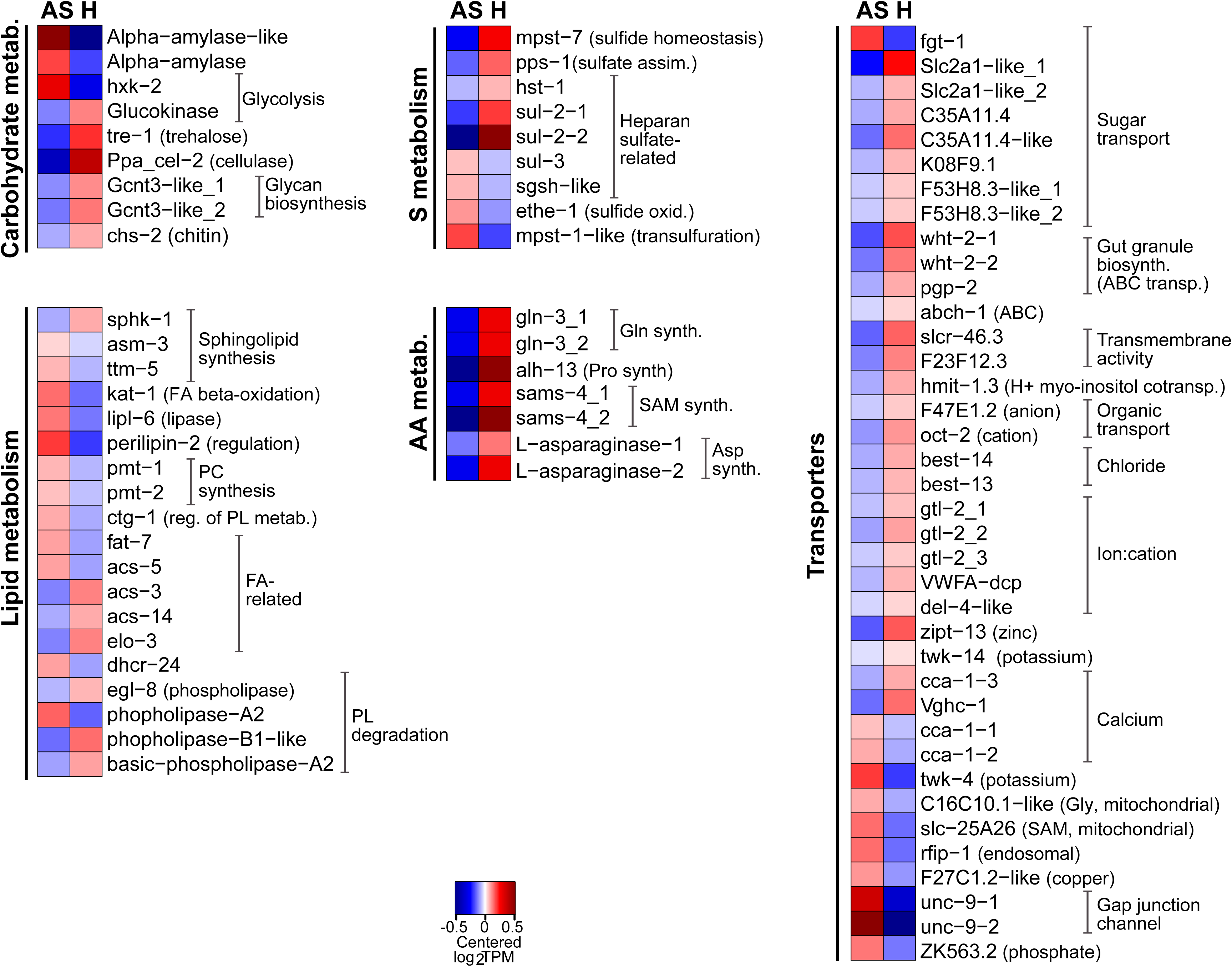
Genes involved in carbohydrate, lipid- and sulfur-metabolism, amino acids biosynthesis, and transport were predominant expressed in hypoxic (H) worms. Heatmap displaying genes upregulated in H relative to AS worms, upon 24 h-long incubations under one of the two conditions (1.5-fold change, FDR ≤ 0.05). Expression levels are displayed as mean-centered log_2_TPM values (transcripts per kilobase million). Genes are ordered by function in their respective metabolic pathways. For each process, the minority of genes that were upregulated in AS worms is shown in Data S1. Red denotes upregulation, and blue downregulation. FA: fatty acids. PC: phosphatidylcholine. PL: phospholipids. Metab: metabolism. Synth: synthesis. Assim: assimilation. Oxid: oxidation. Transp: transporters.

GABA-mediated neurotransmission has been documented for facultative anaerobic animals thriving in anoxic conditions (Lutz et al., 1997; Milton et al., 1998; Lutz et al., 2003a, b). Due to its inhibitory nature, it contributes to avoid membrane depolymerization (Nilsson et al., 1990; Milton et al., 1998). Moreover, given that it relaxes muscles, the increment of GABA may impact the movement of the animal (Mclntire et al., 1993; Schuske et al., 2004). Therefore, upregulation of GABA-mediated neuronal activity might explain why anoxic *L. oneistus* did not form tight worm clusters after 24h (Supplemental movie 3).

#### Dopamine-mediated neurotransmission

A gene encoding for the tyrosine hydroxylase Cat-2 (*cat-2*), which is needed for dopamine biosynthesis (Sawin et al., 2000) and two putative dopamine receptors (*protein-D2-like* and a G_PROTEIN_RECEP_F1_2 domain- containing protein (*dop-5*); Sanyal et al., 2004) were upregulated in AS worms. Moreover, a *dat-1*-like gene mediating dopamine reuptake into the presynaptic terminals was downregulated (Gainetdinov et al., 2002; McDonald et al., 2006) in AS worms (Figure 5).

#### Calcium-binding and -sensing proteins

Finally, in AS worms several calcium-binding or -sensing proteins (e.g., *ncs-2*, *cex-2*, and a calbindin-like (CALB1 homologue); Soontornniyomkij et al., 2012; Hobert et al., 2018; Figure 5), as well as calcium transporters (*cca-1*, Steger et al., 2005; Transport category, Figure 6) were upregulated. On the one hand, we hypothesize their involvement in the inhibitory neural signaling described above (for example, Ncs-2 mediates the cholinergic and GABAergic expression of *C. elegans* (Zhou et al., 2017). On the other, they may protect cells against the stress inflicted by anoxia, which involves calcium overload and consequent cellular acidification (Bickler et al., 1992; Dell’Anna et al., 1996; Galli et al., 2014).

### Genes upregulated in hypoxic (H) nematodes

#### Innate immune pathways and effectors

Animals recognize and respond to microbes by means of immunoreceptors including Toll-like receptors, conserved from sponges to humans (Akira et al., 2006). We identified almost all genes belonging to this pathway, including the one encoding for the NF-kB transcription factor. This came as a surprise given that, up to now, the has not been identified in any other nematode NF-kB (Pujol and Ewbank, submitted). As surprising, was the fact that not only two Toll-like receptors (*tol-1* and *tol-1-like*), but also genes encoding for antimicrobial proteins such as a peroxisome assembly factor involved in defense against Gram- (*prx-11*-like, Wang, D. (2019), a putatively antifungal endochitinase (Dravid et al., 2015) and Bactericidal Permeability Increasing proteins (BPIs) were also more abundant in H worms. BPIs may bind LPS and perforate Gram- membranes and have shown to play a symbiostatic role in other invertebrates (Bruno et al., 2019; Krasity et al., 2015; Chen et al., 2017). However, it is unclear whether activation of the *L. oneistus* Toll pathway leads to the nuclear NF-kB switching on the expression of antimicrobial genes or whether, as shown in *C. elegans*, the Toll pathway mediates behavioral avoidance of pathogens (Pradel et al., 2007; Brandt et al., 2015).

Overall, the apparent oxygen stimulation of a central innate immunity pathway and, directly or indirectly, of broad range anti-defense mechanisms could be adaptations to the fact that in oxygenated environments (when crawling in superficial sand layers), *L. oneistus* is exposed to predation from bigger animals, but also to pathogenic members of the bacterioplankton. Overexpression of broad-range antimicrobials in response to oxygen might therefore help *L. oneistus* to avoid colonization by potentially deleterious, fouling bacteria (e.g., *Vibrios*, *Roseobacters* and *Pseudoaltermonas/Alteromonadales*) when crawling close to the water column (Dang and Lovell, 2016; M. Mussmann, personal communication).

#### Development

Although development-related genes were some of the most expressed under all conditions (Figure 1), many were upregulated in H nematodes (Figure 2 and 5). Among the development-related genes upregulated in H nematodes were those related to molting (e.g., *nas-36*, *nas-38*, *chs-2*, *ptr-5*, *ptr-18*, *apl-1*, *myrf-1*; Suzuki et al., 2004; Zhang et al., 2005; Zugasti et al., 2005; Hornsten et al., 2007; Russel et al., 2011), germ line establishment (e.g., *ccm-3*, *rsks-1*; Pan et al., 2007; Pal et al., 2017), oogenesis/spermatogenesis (*crt-1*, Park et al., 2001), embryonic development and yolk production (*smp-*1, *cpna-1*, *plt-1*, *vit-*6, *crt-1*, *arrd-*17, *mlc-5;* Clark et al., 1997; Goedert et al., 1996; Gatewood et al., 1997; Fuji et al., 2002; Gally et al., 2009; Zahreddine et al., 2010; Jee et al., 2012; Warner et al., 2013; Fisher et al., 2014; Perez and Lehner, 2019), and/or larval development (*nmy-1*, *ifb-1;* Ding et al., 2004; Osório et al., 2019), as well as male tip (Cdt1, *plx-1*, *ver-3*, ; Nelson et al., 2011; Dalpé et al., 2004; Dalpe et al., 2013), vulva morphogenesis (*hda-1*, *unc-62*), and a hermaphrodite-related gene (*hda-1;* Dufourcq et al., 2002; Choy et al., 2007) (Figure 5). Morever, transcripts encoding for a number of proteases shown to be involved in *C. elegans* molting (e.g., *nas-38*, *nas-6*-like; Park et al., 2010), development (e.g., teneurin-a-like; Topf and Drabikoswki, 2019), neuronal regrowth or locomotion (*tep-1*; Kim et al., 2018) and pharingeal pumping (e.g., neprilysin *nep-1*; Spanier et al., 2005) were also more abundant in H worms. Remarkably, *vav-1*, which, besides being involved in male tip and vulva morphogenesis (Nelson et al., 2011), may also regulate the concentration of intracellular calcium (Norman et al., 2005), was one of the few development-related genes to be downregulated in H nematodes (see previous section on Ca-binding proteins).

To sum up, and as expected, the host appears to exploit oxygen availability to undertake energetically costly processes, such as development and molting (De Cuyper and Vanfleteren 1982; Uppaluri and Brangwynne 2015).

#### Carbohydrate metabolism

If in AS nematodes, glycogen or starch appeared prominent carbon sources, H worms seemed to exploit trehalose and cellulose instead. Indeed, genes that degrade trehalose (*tre-1*, Pellerone et al., 2003) and cellulose (Ppa-*cel-2*, Schuster et al., 2012) were upregulated in H worms, as well as a putative ADP-dependent glucokinase (C50D2.7) involved in glycolysis (Yuan et al., 2012). The use of this pathway was supported by the overexpression of four genes encoding for sugar transporters (Slc2-A1, C35A11, K08F9.1, F53H8.3; Kitaoka et al., 2013; Bertoli et al., 2015), perhaps switched on by active mTOR (see above) (Figure 6) (Howell et al., 2011).

Additionally, *L. oneistus* appeared to exploit oxygen to synthesize complex polysaccharides, such as heparan sulfate (*hst-1-*like; Miyagawa et al. 1988; Bhattacharya et al., 2009) and glycan (Gcnt3-like) (Figure 6), as an ortholog of the N-deactetylase/N- sulfotransferase *hst-1*, related to heparin biosynthesis was also upregulated (Bhattacharya et al., 2009).

Although glycolysis seems to generate ATP in both AS and H worms, it is not clear why the latter would prefer to respire cellulose or trehalose instead of starch. Given its role as a membrane stabilizer, we speculate that AS worms might prioritize the storage of trehalose over its degradation to preserve membrane integrity (Figure 6) (Crowe et al 1987; Carpenter et al., 1988; Clegg et al., 1997; Chen et al., 2002; Haddad 2006). Of note, based on its genome draft, the symbiont may synthetize and transport trehalose, but it may not use it (Paredes et al., 2021). Therefore, we hypothesize symbiont-to-host transfer of trehalose under hypoxia. Consistently, the symbiont’s trehalose synthesis-related gene (*otsB*; Paredes et al., 2021), and the host trehalase (*tre-1*; Figure 6) were both upregulated under hypoxia and metabolomics could detect trehalose in both partners (Table S1). Metabolomics also detected sucrose in both the holobiont and the symbiont fraction (Table S1). Given that, based on transcriptomics and proteomics, the nematode can utilize sucrose but cannot synthesize it (Data S1), whereas the symbiont can (Paredes et al., 2021), as in the case for trehalose, we hypothesize symbiont-to-host sucrose transfer.

#### Acetylcholine-mediated neurotransmission

Instead of upregulating genes involved in inhibitory (GABA and dopamine-mediated) neurotransmission, hypoxic worms appeared to use excitatory acetylcholine-mediated neurotransmission as indicated by the upregulation of *molo-1*, *acr-20,* cup-4, *lev-9*, and sphingosine kinase *sphk-1* that promotes its release (Mongan et al., 2002; Patton et al., 2005; Gendrel et al., 2009; Boulin et al., 2012; Chan et al., 2012) (Figure 5). On the one hand, acetylcholine-mediated neurotransmission might promote ROS detoxification in H worms (Sun et al., 2014). On the other hand, its downregulation in AS worms may beneficially decrease calcium influx (Hochachka and Lutz, 2001).

#### Feeding, mating, mechanosensory behavior and axon guidance and fasciculation

Transcripts related to the neuronal regulation of energy-demanding activities such as feeding, mating, motion, as well as nervous system development were more abundant in H nematodes (Figure 5, and Data S1). More precisely, upregulated genes were involved in pharyngeal pumping (*nep-1*, *lat-2*; Spanier et al., 2005; Guest et al., 2007), male mating behavior and touch (*pdfr-1*, *tbb-4, ebax-1,* Hurd et al., 2010; Wang, Z. et al., 2013), axon guidance and fasciculation (*spon-1*, *igcm-1*, *ebax-1*, *tep-1*; Kim et al., 2018; Woo et al., 2008; Schwarz et al., 2009; Wang, Z et al., 2013), mechanosensory behavior (e.g., *mec-12*, *delm-2*; Gu et al., 1996; Han et al., 2013). Additionally, we also observed the upregulation of a gene encoding for a glutamate receptor (*glr-7*) possibly involved in feeding facilitation (Li et al., 2012).

#### Amino acid biosynthesis

Transcripts of genes involved in the synthesis of glutamine and proline (*gln-3* and *alh-13*, respectively), aspartate (L-asparaginases; Tsuji et al., 1999) and S-adenosyl-L-methionine (SAM) (*sams-4*; Chen et al., 2020) were all upregulated in H worms (Figure 6), as well as one encoding for the ornithine decarboxylase *odc-1* which is involved in biosynthesis of the polyamine putrescin, and is essential for cell proliferation and tissue growth (Russell et al., 1968; Heby, 1981). Moreover, polyamines, with their high charge-to-mass ratio may protect against superoxide radicals, which, as mentioned, harm cell membranes and organelles, oxidize proteins, and damage DNA (Gilad et al., 1991; Longo et al., 1993).

#### Lipid biosynthesis

Genes upregulated in H worms mediate the biosynthesis of long chain fatty acids (*acs-3*, *acs-14*, *elo-*3 but not *acs-5*; Yuan et al., 2012; Ward et al., 2014; Wang et al., 2021), sphingolipids (a sphingosine kinase-1 (*sphk-1*) and *egl-8*, which controls egg laying and pharyngeal pumping in *C. elegans* (Bastiani et al., 2003). Notably, sphingolipids may be anti-apoptotic (Taha et al., 2006) or result in acetylcholine release (Chan et al., 2012).

On the other hand, ceramides, which have antiproliferative properties and who may mediate resistance to severe oxygen deprivation (Deng et al., 2008; Menuz, et al. 2009), appeared to be mainly synthesized in AS worms, as indicated by the upregulation of genes involved in ceramide biosynthesis (*asm-3*, *ttm-5*; Watts et al., 2017) (Figure 6).

#### Transport

As anticipated in the introduction, anoxia-tolerant animals switch off ATP-demanding processes such as ion pumping (Lutz et al., 1996; Galli et al., 2014). Indeed, transcripts encoding for proteins involved in cation channel activity (*gtl-2*, voltage gated H channel 1; Teramoto et al., 2010), sodium transport (*delm-2*-like; Han et al., 2013), chloride transport (*anoh-1*, *best-13*, *best-14*; Tsunenari et al., 2013; Wang, Y. et al., 2013; Goh et al., 2018), ABC transport (*wht-2*, *pgp-2*, slcr-46.3, F23F12.3, *hmit-1.3;* Currie et al., 2007; Schroeder et al., 2007; Kage-Nakadai et al., 2011) and organic transport (F47E1.2, *oct-2*; Pao et al., 1998) were all more abundant in H than AS worms (Figure 6).

#### Sulfur metabolism

The *mpst-7* gene which is involved in organismal response to selenium and it is switched on in hypoxic *C. elegans* (Romanelli-Credrez et al., 2020) was upregulated in H nematodes (Figure 6). Given that the latter is thought to catalyze the conversion of sulfite and glutathione persulfide (GSSH) to thiosulfate and glutathione (GSH) (Filipovic et al., 2018), hypoxia-experiencing *L. oneistus* might express this enzyme to recharge the cells with GSH and hence, help to cope with oxidative stress (Hayes and McLellan, 1999; Mytilineou et al., 2002; Diaz-Vivancos et al., 2015). Also more abundant in H worms were transcripts encoding for the sulfatases 2 (*sul-*2) (Morimoto-Tomita et al., 2002) and a PAPS-producing *pps-1* (3 -phospho-adenosine-5 -phosphosulfate (PAPS) considered the universal sulfur donor; Bhattacharya et al., 2009), as well as for the chaperones *pdi-6* and protein-disulfide-isomerase-A5-like which require oxygen to mediate correct disulfide bond formation in protein folding (Teodoro and O’Farrell, 2003; Rose et al., 2017; Livshits et al., 2017) (Figure 6).

Conversely, a putative sulfide-producing enzyme (*mpst-1*) who protects *C. elegans* from mitochondrial damage (Qabazard et al., 2014; Ng et al., 2019; Kimura, 2020) was upregulated in AS nematodes. Notably, under AS, *L. oneistus* might detoxify sulfide by producing glutathione and taurine (Rose et al., 2017), as a persulfide dioxygenase (*ethe-1*) and a cysteine dioxygenase (*cdo-1*) which catalyzes taurine synthesis via cysteine degradation were upregulated. Sulfide detoxification via taurine accumulation is a common strategy in chemosynthetic animals (reviewed in Cavanaugh et al., 2006).

All in all, *L. oneistus* appeared to limit excess accumulation of free sulfide in anoxia and to free sulfate when oxygen was available.

## Conclusions

Overall and irrespectively of the conditions it was subjected to, *L. oneistus* mostly expressed genes involved in degradation, energy generation, stress response and immune defense. Astonishingly, *L. oneistus* did not enter suspended animation when subjected to anoxic sulfidic conditions for days. We hypothesize that in the absence of oxygen, ATP production is supported by trehalose and cellulose catabolism, and by rewiring the ETC in such way as to use rhodoquinone (RQ) as electron carrier, and fumarate as electron acceptor. Moreover, the nematode activates several degradation pathways (e.g., ubiquitin-proteasome system (UPS), autophagy, and apoptosis) to gain nutrients from anoxia- or ROS-damaged proteins and mitochondria. Further, AS worms also upregulated genes encoding for ribosomal proteins and putative symbiont-binding proteins (lectins). Finally, as proposed for other anoxic- tolerant animals, the worm seems to upregulate its antioxidant capacity in anticipation of reoxygenation. When in hypoxic conditions (Figure 7, left), instead, we speculate that the worm uses starch for energy generation to engage in costly developmental processes such as molting, feeding, and mating, likely relying on excitatory neurotransmitters (e.g., acetylcholine), and it upregulates the Toll immune pathway and, directly or indirectly, the synthesis of broad range antimicrobials (e.g., fungicides, bactericidal permeability increasing proteins).

**Figure 7.**
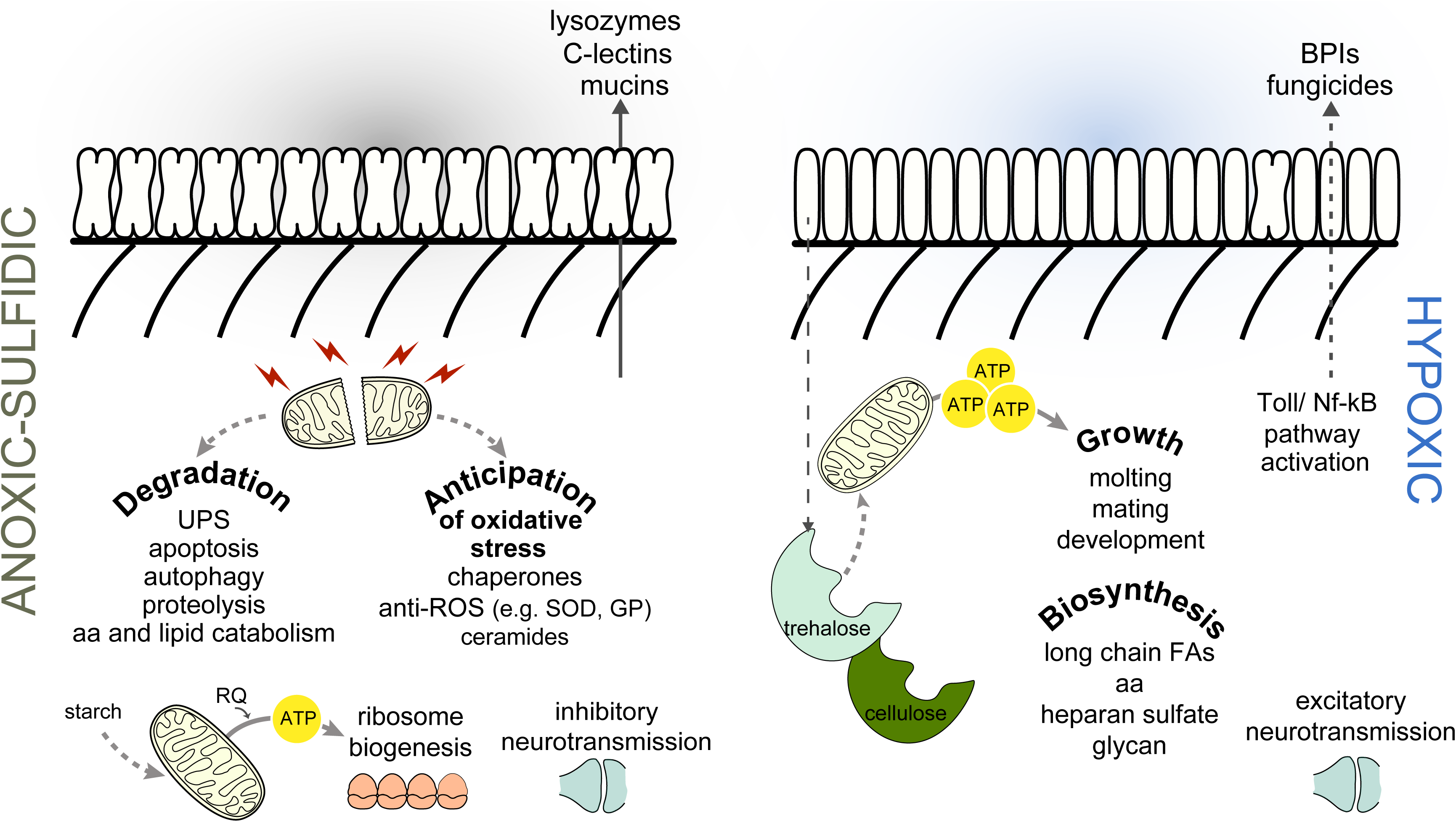
Schematic representation of *Laxus oneistus* physiology in anoxic and hypoxic sand. In anoxic sulfidic sand (left) *L. oneistus* does not enter suspended animation. Instead, it upregulates the expression of genes mediating inhibitory neurotransmission, involved in symbiosis establishment (e.g., lectins, mucins) and in ribosome biogenesis. Metabolism may be supported by the degradation of starch and by rewiring the electron transfer chain: rhodoquinone (RQ) is used as electron carrier and fumarate as electron acceptor. Moreover, the worm activates degradation pathways (e.g., ubiquitin-proteasome system (UPS), autophagy, and apoptosis) and may anticipate reoxygenation by upregulating superoxide dismutase (SOD) and glutathione peroxidase (GP). In hypoxic sand (right), instead, *L. oneistus* appears to use trehalose and cellulose for energy generation, while engaging in costly processes such as development, molting, feeding, and mating. Genes involved in excitatory neurotransmission are also upregulated, together with Toll receptors and immune effectors (e.g., fungicides, bactericidal permeability increasing proteins).

When looking at the *Laxus*-*Thiosymbion* symbiosis in light of what was recently published (Paredes et al., 2021), we could identify two signs of inter-partner metabolic dependence: in anoxia worms might transfer lipids to their symbionts, and in hypoxia the symbionts might transfer trehalose to their hosts.

Furthermore, we may conclude that, wherever in the sand the consortium is, one of the two partners is bound to be stressed: in anoxia, the symbiont appear to proliferate more, while its animal host engages in degradation of damaged proteins and mitochondria and in detoxification. In the presence of oxygen, the situation is inverted: the symbiont seems massively stressed, while the host can afford energy costly biosynthetic processes to develop and reproduce (Figure 7). It is therefore fascinating that, in spite of the dramatically different needs a bacterium and animal must have, the *Laxus*-*Thiosymbion* symbiosis evolved.

## Supporting information

Suppl. Movie 1 (anoxic 6d)

Supp. Movie 2 (anoxic T0)

Suppl. Movie 3 (anoxic T24)

Suppl. Movie 4 (oxic T24)

## ACKNOWLEDGEMENTS

This work was supported by the Austrian Science Fund (FWF) grant P28743 (T.V., S.B., and L.K.), the FWF DK plus grant W1257: Microbial Nitrogen Cycling (G.F.P., L.K.), the FWF DOC 69 doc.fund (T.V). We thank Yin Chen for providing the facilities for lipidomics analysis, and Marvin Weinhold’s, Jana Matulla’s and Sebastian Grund’s excellent technical work during metabolite analysis, and protein sample preparation and MS analysis, respectively. We are grateful to the Carrie Bow Cay Marine Field Station, Caribbean Coral Reef Ecosystem Program, and Station Manager Zach Foltz and Scott Taylor for their continuous support during field work. We thank Nicole Dubilier for access to data on *Olavius algarvensis,* and Jonathan Ewbank and Marc Mussmann for insightful comments on the manuscript. Finally, we were inspired by insightful discussions with Monika Bright and Jo□rg A. Ott. This is contribution number XXX of the Carrie Bow Cay Marine Field Station, Caribbean Coral Reef Ecosystem Program.

## MATERIALS AND METHODS

### Sample collection

*Laxus oneistus* individuals were collected on multiple field trips (2016-2019) at approximately 1 m depth from sand bars off the Smithsonian Field Station, Carrie Bow Cay in Belize (16°48’11.01’’N, 88°4’54.42’’W). The collection of the nematodes, the incubations set up for RNA sequencing, lipidomics, proteomics and metabolomics, as well as the RNA extraction, and library preparation are described in Paredes et al., 2021. Importantly, the nematodes had a bright white appearance and replicate incubations were started simultaneously. Note that the Supplemental material describes changes in the lipidomics and proteomics pipelines, as well as the metabolomics, and sequencing data of *Olavius algarvensis*.

### Host transcriptome de novo assembly

In preparation for the assembly, reads from each sample were first mapped to the symbiont as described before (Paredes et al., 2021), and remaining rRNA reads from all domains of life were removed from unmapped reads using sortmerna v2.1 in combination with the SSURef_NR99_119_SILVA_14_07_14 and LSURef_119_SILVA_15_07_14 databases. Further, exact duplicate reads were removed using PRINSEQ lite’s derep option. Read files free of symbiont reads, rRNA reads and exact duplicates were used as input for transcriptome sub-assemblies via Trinity v2.6.6 with the strand-specific option (--SS_lib_type F) (Grabherr et al., 2011). Two sub-assemblies differing in the number and type of input read files were performed: (1) 9 input read files including biological triplicates from 3 incubation conditions (O, H, A) and (2) 4 input read files including a single replicate from 4 incubation conditions (O, H, A and hyper-O). Hyper-O refers to an incubation in which air was pumped directly into the exetainers for the entire incubation period to supersaturate the seawater (300 %O_2_). However, as this incubation condition yielded an incongruous transcriptional response by the symbiont (data not shown), these read data were only used to extend the host transcriptome’s coding repertoire. The qualities of both sub-assemblies were assessed as described below.

We then performed an intra-assembly clustering step as described in (Cerveau and Jackson, 2016), during which identical transcripts were removed from the sub-assemblies using CD-HIT-EST (Fu et al., 2012). To further reduce redundant transcripts, only the longest isoform for each ‘gene’ identified by Trinity was kept using Trinity’s get_longest_isoform_seq_per_trinity_gene.pl utility. The remaining transcripts of each sub- assembly were then concatenated to produce a merged transcriptome assembly. The final assembly was created by applying another sequence clustering using CD-HIT-EST to avoid inter-assembly redundancy. Here, the identity parameter of 80% (-c 0.8) combined with a minimal coverage ratio of the shorter sequence of 80% (-aS 0.8) and minimal coverage ratio of the longest sequence of 0.005% (-aL 0.005) yielded the best-performing assembly in terms of number of transcripts (162,455) and contiguity (N50 value of 770) (data not shown).

Assembly completeness was assessed by estimating completeness via BUSCO nematode single-copy orthologs (Simão et al., 2015). Importantly, the merged assembly yielded a higher BUSCO-based completeness compared with the two sub-assemblies; 79.2% of the BUSCO nematode single-copy orthologs were found to be present and complete in the final assembly (636 single-copy/142 duplicated), whereas assembly (1) scored 77.8% (233 single-copy/531 duplicated) and assembly (2) was 76.2% complete (314 single-copy/434 duplicated). Further, assembled transcripts were filtered based on taxonomic classification. Transcripts were matched against the RefSeq protein database using blastx (E value 1E-3), and the output was then used as input for taxonomic assignment via MEGAN v5 (Huson et al., 2007). Only transcripts classified as belonging to ‘Eukarya’ were kept (MEGAN parameters: Min Score: 50, Max Expected: 1E-2, Top Percent: 2), which reduced the number of putative *L. oneistus* transcripts to 30,562. Assembled transcripts were also functionally annotated using Trinotate (Bryant et al., 2017). Briefly, predicted protein coding regions were extracted using TransDecoder (https://github.com/TransDecoder), both transcripts and predicted protein sequences were searched for protein homology via blastx and blastp, respectively, and predicted protein sequences were annotated for protein domains (hmmscan), signal peptides (signalP) and transmembrane domains (THMMM). 85,859 transcripts exhibited at least one functional annotation. Finally, only taxonomy-filtered transcripts with at least one functional annotation were kept, thereby further reducing the number of putative host transcripts to 27,984, with 22,072 thereof predicted to contain protein coding regions. BUSCO-based completeness for this filtered host transcriptome assembly was 78.8% (635 single-copy/139 duplicated).

### Gene expression analysis

Raw sequencing reads quality assessment and preprocessing of data was followed as described in Paredes et al., 2021. Trimmed reads were mapped to the de novo transcriptome assembly and transcript abundance was estimated using RSEM v1.3.1 (Li and Dewey 2011) in combination with bowtie with default settings except for the application of strandedness (-- strandedness forward). Read counts per transcript were used for differential expression analysis, and TPM (transcripts per kilobase million) values were transformed to log2TPMs as described in Paredes et al. 2021.

Gene and differential expression analyses were conducted using the R software environment and the Bioconductor package edgeR v3.28.1 (Gentleman et al., 2004; Robinson et al., 2010; R core Team, 2013), and as shown in Paredes et al., 2021. Here, we only describe the modifications that were made to the pipeline. Genes were considered expressed if at least ten reads in at least three replicates of one of the four conditions could be assigned. Excluding the replicates of the oxic condition, we found that 74.9% of all predicted nematode protein-encoding genes to be expressed (16,526 genes out of 22,072). Log_2_TPM were used to assess sample similarities via multidimensional scaling based on Euclidean distances (R Stats package) (R core Team, 2013) (Figure S1B), and the average of replicate log_2_TPM values per expressed gene and condition was used to estimate expression strength. Median gene expression of entire metabolic processes and pathways per condition was determined from average log_2_TPM values.

Expression of genes was considered significantly different if their expression changed 1.5-fold between two treatments with a false-discovery rate (FDR) ≤ 0.05 (Rapaport et al., 2013). Throughout the paper, all genes meeting these thresholds are either termed differentially expressed or up- or downregulated. For the differential expression analyses between the AS, H and A conditions see Data S1. Heatmaps show mean-centered log_2_TPM expression values to highlight gene expression change.

All predicted *L. oneistus* proteins were automatically annotated using eggNOG-mapper v2 (Cantalapiedra et al., 2021) against eggNOG 5.0 (Huerta-Cepas et al., 2019) using diamond v2.0.4 (Buchfink et al., 2021). All genes that are shown and involved in a particular process were manually curated by blasting them against both the NCBI BLASTP nr database (Altschul et al., 1990) and the WormBase (Harris et al., 2020; https://wormbase.org/tools/blast_blat).

## Data availability

This Transcriptome Shotgun Assembly project has been deposited at DDBJ/EMBL/GenBank under the accession GJNO00000000. The version described in this paper is the first version, GJNO01000000. RNA-Seq data are available at the Gene Expression Omnibus (GEO) database and are accessible through accession number GSE188619.

## SUPPLEMENTAL MATERIAL LEGENDS

**Figure S1. Experimental conditions, sample similarity and differential expression.** (A) Experimental setup was previously described (Paredes et al. 2021). Briefly, nematodes were subjected to different oxygen concentrations for 24 h: anoxic with sulfide (AS: 0mM O_2_, 25mM sodium sulfide added), anoxic without sulfide (A, 0mM O_2_), hypoxic (H, 60mM O_2_ after 24 h), and oxic (O, 100mM O_2_ after 24 h). The box around the anoxic incubation vials illustrates that these incubations were carried out in a polyethylene glove bag. (B) Similarity between transcriptome samples based on Euclidean distances between expression values (log_2_TPM), and visualized by means of multidimensional scaling (C) Differential gene expression (DE) analysis between incubations showed that the number of DE genes was low (maximum value was 4.8% of all expressed genes for the H vs AS conditions). Genes were considered differentially expressed if their expression changed 1.5-fold with a false-discovery rate (FDR) of ≤ 0.05.

**Figure S2.** Statistical analysis, relative transcript abundance and expression levels of the top 100 detected proteins of *L. oneistus* across all conditions. (A) Relative protein abundance (%) of the top 100 detected proteins present in a particular manually curated functional category. The top 100 proteins were collected by averaging the expression values across all replicates of all incubations (Figure S1A, Data S2). Functional classifications were extracted from the curated database UniProt and from comprehensive literature search focused mainly on *C. elegans,* and confirmed with the automatic annotated eggNOG classification (Data S1). (B) Median gene expression levels of selected *L. oneistus* manually annotated functional categories of the top 100 expressed proteins. Each dot represents the average %cOrgNSAF per protein across all replicates of all incubations. Notice that some categories were created with genes of overlapping functions (e.g., cytoskeleton/locomotion/nervous system). All protein names (or locus tags for unidentified protein names) are listed in Data S2.

**Figure S3. Transcriptomics vs proteomics comparison.** Pearson correlation between all transcripts and proteins (Data S1) automatically classified based on their functional category. The Pearson correlation between all expressed transcripts and all detected proteins (r = 0.4) was found to be low (Figure S3).

**Figure S4. Relative transcript abundance and expression levels of the top 100 expressed genes of *O. algarvensis* across all conditions.** (A) Relative transcript abundance (%) of the top 100 expressed genes with a manually curated functional category. The top 100 expressed genes were collected by averaging the expression values (log_2_TPM) across all replicates of all incubations (see Supplemental material). Functional classifications were extracted from the curated database UniProt and from comprehensive literature search focused mainly on *C. elegans*). (B) Median gene expression levels of selected *O*. *algarvensis* manually annotated functional categories of the top 100 expressed genes. Metabolic processes include both differentially and constitutively expressed genes. Each dot represents the average log2TPM value per gene across all replicates of all incubations.

**Figure S5. *L. oneistus* lipid composition in anoxic and oxic conditions after 24 h.** Major lipid classes and their abundance relative to all lipids detected showed no statistical difference between both conditions. For details on methodology see Supplemental material.

**Table S1.** Metabolites detected in at least two biological replicates of either the holobiont fraction (*Laxus oneistus* and its ectosymbiont) or in the symbiont fraction (see Supplemental material). RT: retention time. Area: area of a peak from a specific compound detected in the GC-MS chromatograms. Grey boxes: no metabolites detected. Blank boxes: Unknown metabolites that are either below the detected threshold (< 700) or might be products of derivatization reagents. Note that cholestane and ribitol were used as internal standards.

**Data S1.** *Ca*. T. oneisti genes, functional annotations, transcript and protein expression.

**Data S2**. Top 100 expressed genes (RNA-Seq) and detected proteins (proteomics data).

**Supplemental video 1.** A batch of 50 *Laxus oneistus* after 6 days in anoxic seawater.

**Supplemental video 2.** A batch of 50 *Laxus oneistus* at the beginning (T0) of the incubations.

**Supplemental video 3.** A batch of 50 *Laxus oneistus* after 1 day (T24 h) in anoxic sulfidic seawater (0 % air saturation, 25 µM H_2_S).

**Supplemental video 4.** A batch of 50 *Laxus oneistus* after 1 day (T24 h) in oxic seawater (87 % air saturation, 0 µM H_2_S).

