## Supplementary figures and images for "Differential regulation of degradation and immune pathways underlies adaptation of the ectosymbiotic nematode *Laxus oneistus* to oxic-anoxic interfaces"

### Figure S1

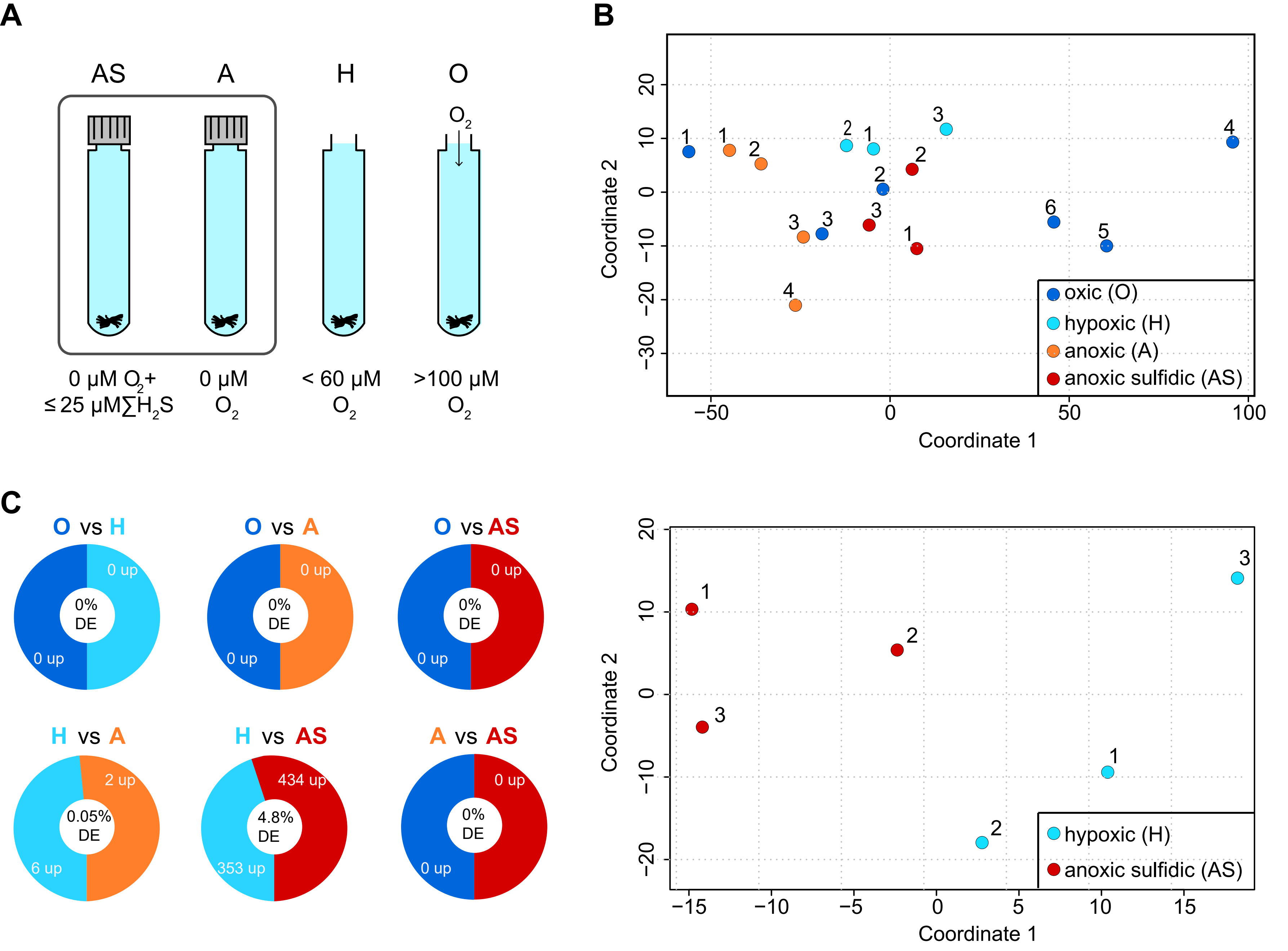

### Figure S2

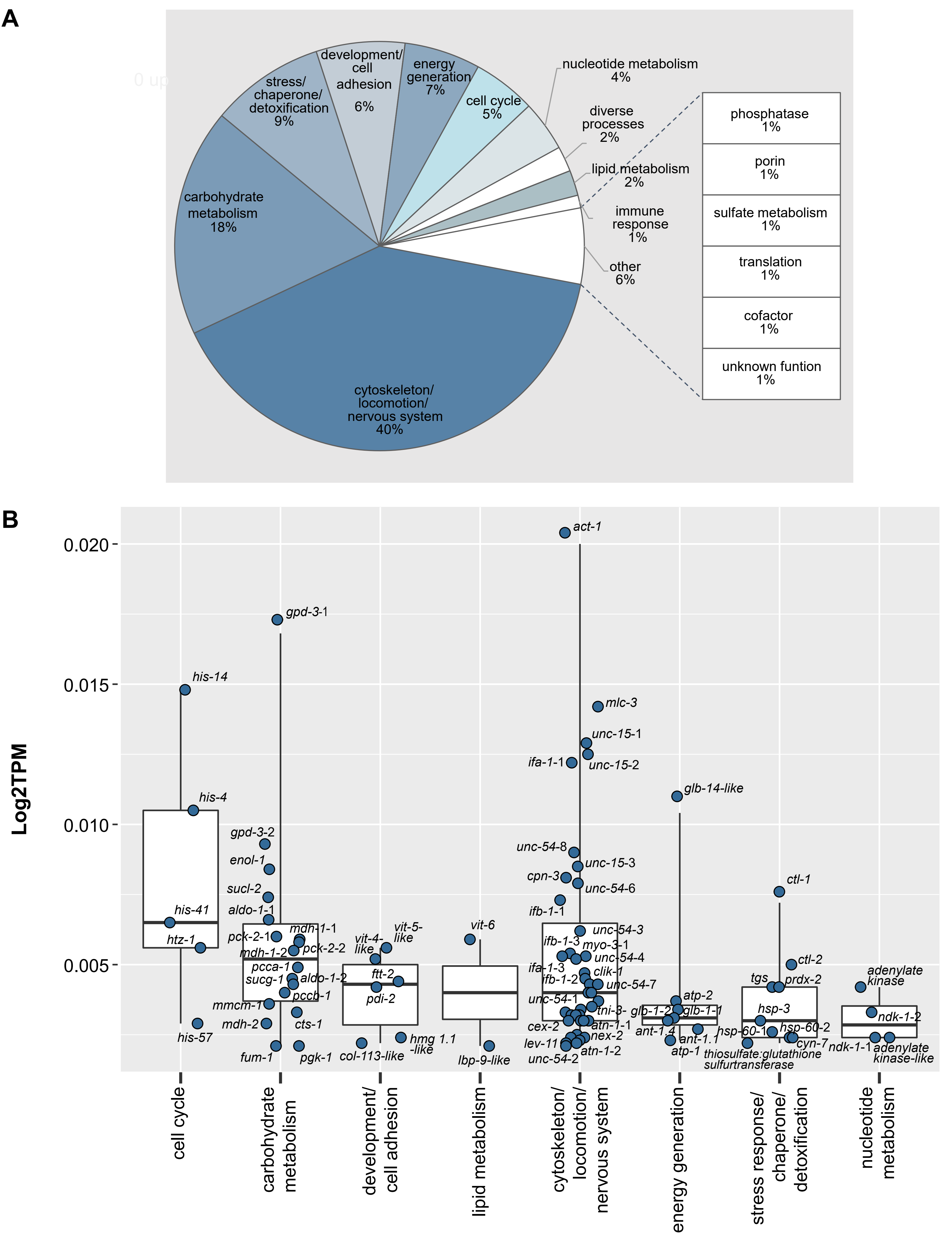

### Figure S3

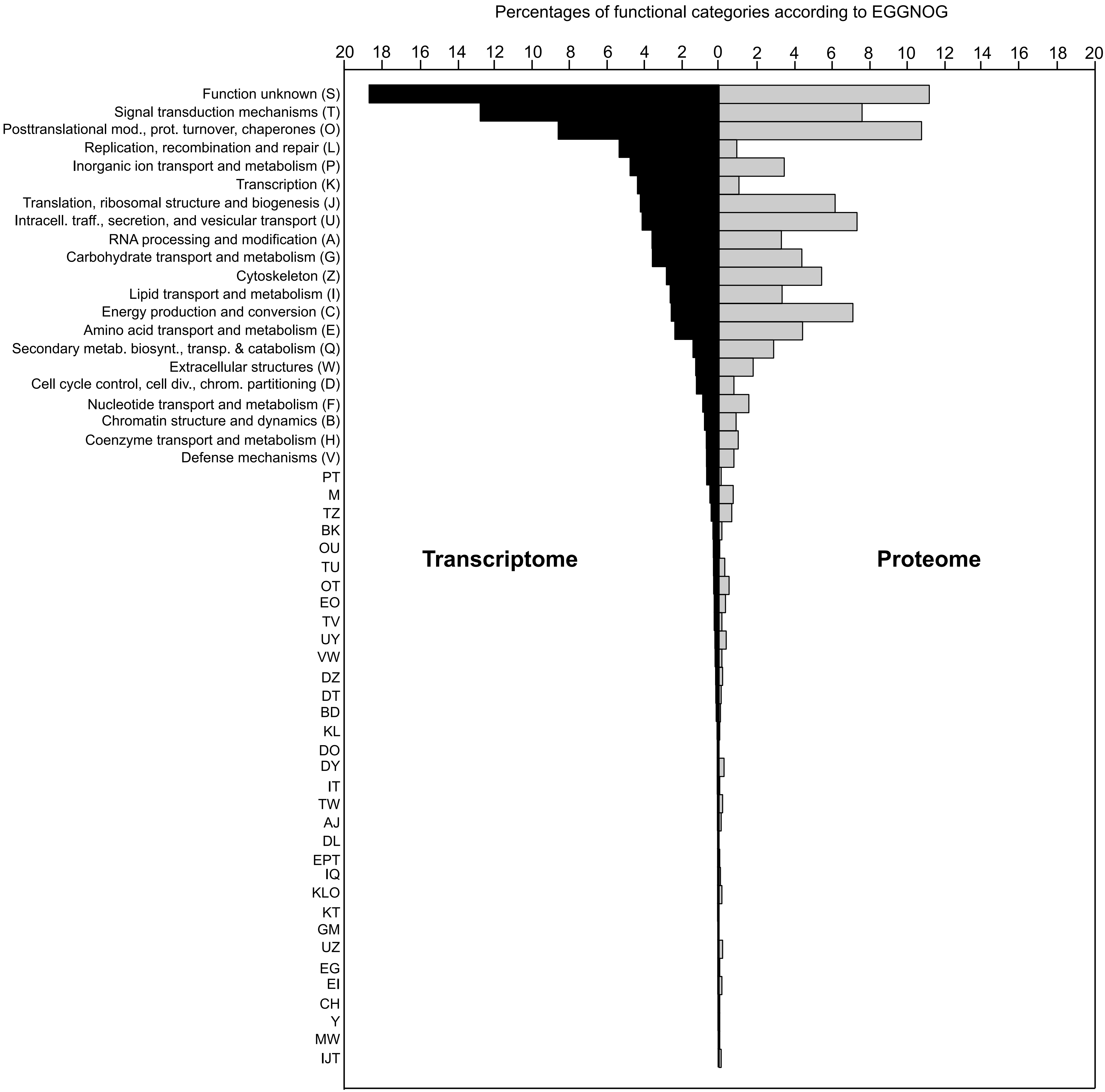

### Figure S4

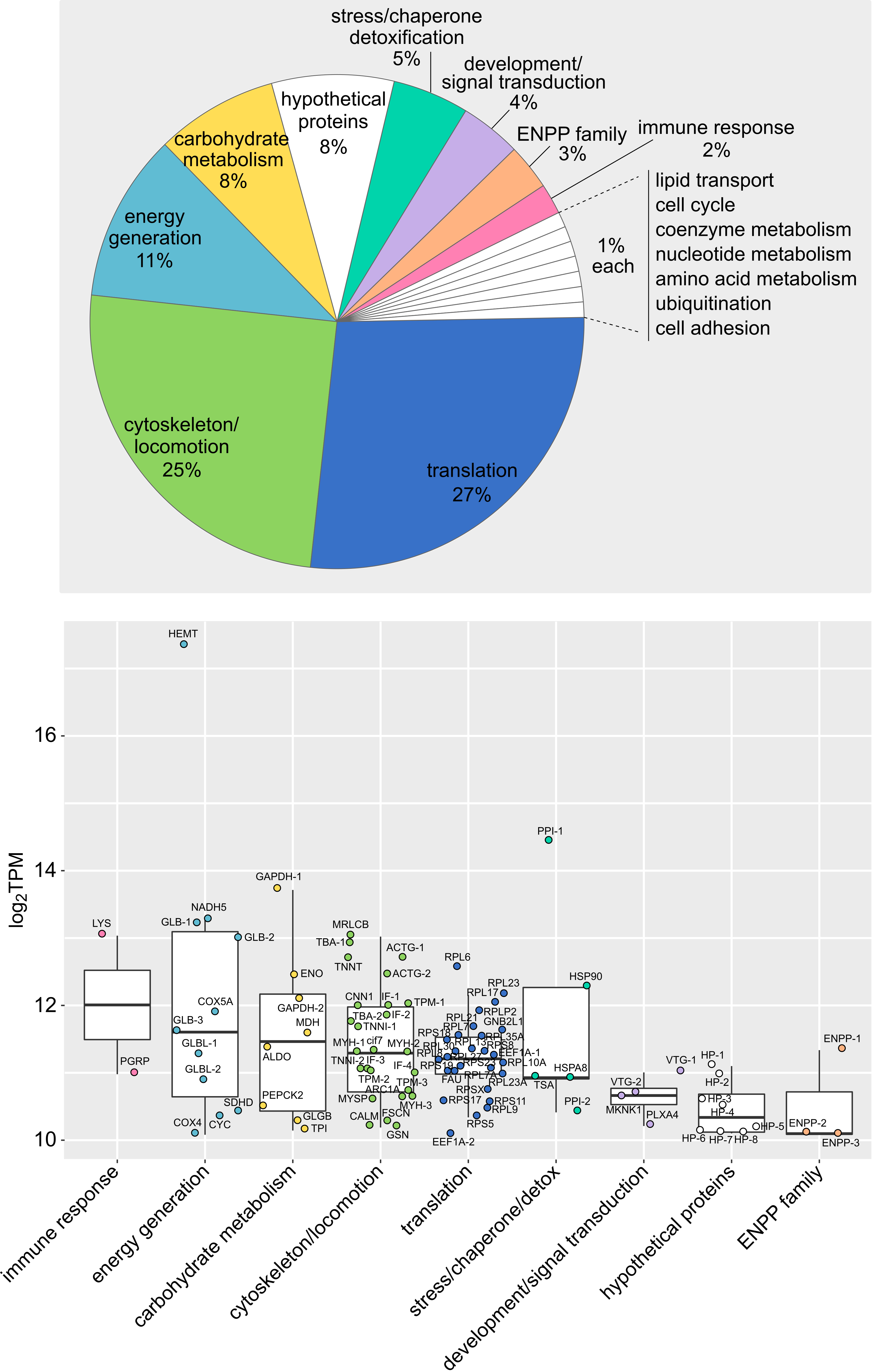

### Figure S5

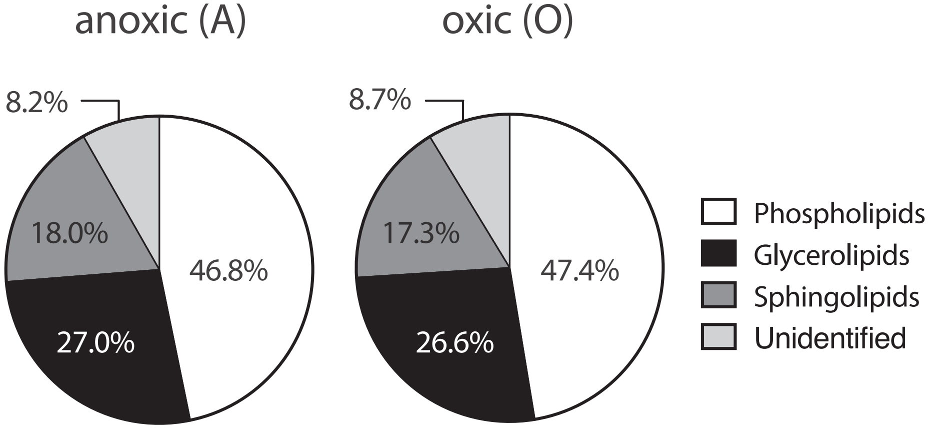
