## Supplementary material for "Differential regulation of degradation and immune pathways underlies adaptation of the ectosymbiotic nematode *Laxus oneistus* to oxic-anoxic interfaces": Suppl. Material & Methods

**Supplemental Materials and Methods**

Paredes et al.

***Olavius algarvensis* transcriptome analysis.** For transcriptomic analyses of *O. algarvensis*, RNA sequences were obtained from 12 individuals collected from Sant’ Andrea bay off the island of Elba (42°48'31"N / 10°08'33"E). Six animals were incubated in anoxic artificial seawater in gas-tight serum bottles for 24 hours before fixed in RNAlater (anoxic treatment). Separate six animals were incubated first in anoxic artificial seawater for 24 hours and subsequently incubated in oxygenated artificial seawater for 12 hours before fixation (oxic treatment). RNA was separately extracted from the 12 individuals with the AllPrep DNA/RNA kit (Qiagen), following the default protocol with a 3 min bead-beating step at 20 Hz. Their cDNA was synthesized with the Ovation RNA-Seq System (NuGEN Technologies Inc., Redwood City, CA) and sheared with the Covaris untrasonicator (Covaris, Woburn, MA). Sequencing libraries were prepared with the NEBNext Ultra DNA Library Prep Kit for Illumina (New England Biolabs, Ipswich, MA), targeting the insert size of 250 bp. Paired-end metatranscriptomic reads of approximately 6 giga bases (100 bases x 2; 30 million read pairs) per sample were generated, using the HiSeq2500 System (Illumina) at the Max-Planck-Genome-Centre Cologne.

A transcriptomic reference of *O. algarvensis* was generated by *de novo* co-assembly of the metatranscriptomes from the 12 individuals. Raw sequences were first de-contaminated from residual Illumina adapters and PhiX sequences and quality-filtered, using bbduk (BBTools; https://jgi.doe.gov/data-and-tools/bbtools/). Sequencing errors were corrected using SEECER (Le et al. 2013). Sequences of *O. algarvensis* mitochondrial genome (mtDNA) and symbiotic bacteria, any ribosomal RNA (rRNA) and DNA contaminants (e.g., *Drosophila sp*.) were removed using bbmap (BBTools). Mapping references were mtDNA and symbiont genomes in *O. algarvensis* (Woyke *et al*. 2006, Sato *et al*. 2020), rRNA sequences from sortMeRNA database (Kopylova et al. 2012), and the *Drosophila melanogaster* genome (GenBank assembly accession: GCA_000001215.4). Reads were assembled using Trinity v2.4.0 (Grabherr et al. 2011). Contigs shorter than 600 bases and with less than 4× coverage were removed to minimize the assembly artifact. Contigs without open reading frames (ORFs) were further removed using TransDecoder v3.0.1 (Haas et al. 2013). To remove contaminating viral, bacterial, archaeal and contaminating eukaryotic (e.g. Cnidaria, Viridiplantae) sequences, taxonomic affiliations of contigs were identified using DIAMOND blastx search v0.8.36.98 (Buchfink et al. 2015) against the NCBI-nr protein database (https://www.ncbi.nlm.nih.gov/; accessed 2018 February), and contigs assigned to Bilateria were exported using MEGAN v6.19.6 (Huson et al. 2007). Redundant contigs due to isoforms and allelic variations were removed with Corset v1.06 (Davidson and Oshlack 2014), with the anoxic vs. oxic treatments used as a grouping factor to differentiate paralogs. Only the longest contigs per gene were kept based on Trinity isoform tags. Completeness of the final transcriptomic assembly was scored at 73.9% with BUSCO v2.01 (Simão et al. 2015) using the metazoan single copy orthologous gene database (odb9). Functional annotation was performed with Blast2GO using OmicsBox v1.4.12 (BioBam Bioinformatics, Valencia) with NBCI-blast against the nr_v5 database and InterPro scan against GO mapping v2019.06.
For gene expression analyses of the 12 *O. algarvensis* individuals treated with the anoxic and oxic treatments above (n = 6 each), the quality controlled sequences were mapped against the transcriptomic reference using RSEM v1.3.1 (Li and Dewey 2011). The TPM values were analyzed as described in the main document. As gene expression profiles were not clustered based on the treatment (data not shown), the log_2_TMP values were averaged across all the individuals, and the top 100 expressed gene categories were identified.

**Proteomics on *L. oneistus*.** Sample collection and preparation of *L. oneistus* pellets for proteomics, as well as the proteins extraction procedure followed by LC-MS/MS analysis were described previously (Paredes et al., 2021)

**Protein identification and quantification***.* The database for protein identification was constructed as described previously (Paredes et al., 2021), but with optimized host sequences derived from host transcriptome *de novo* assembly (see main text), and contained 21,721 *Laxus oneistus* host proteins, 5,145 *Ca*. T. oneisti proteins (JAAEFD000000000), and a set of 42 common laboratory contaminants. In this way, 2,626 host proteins and 1,348 symbiont proteins were identified in total. Data S1 indicates all identified host proteins in the column “Proteome detection” (Column AB). Relative abundance of identified proteins was calculated from total spectrum counts as normalized spectral abundance factor (%NSAF) values – giving the percentage of each protein relative to all proteins in the respective sample (Florens et al., 2006), and as %OrgNSAF, giving a protein’s percentage relative to all host proteins in the respective sample (Mueller et al., 2010). %OrgNSAF values are listed in Data S1 (columns AI and AJ).

**Intact polar lipid extraction and analysis.** Sample collection and preparation of *L. oneistus* pellets for lipidomics, as well as lipid extraction from the nematodes were described previously (Paredes et al., 2021). Modifications on the latter are described next.

Briefly, pelleted nematodes were taken up in 1.5 ml 0.2 µm filtered seawater and 0.5 ml were transferred to 2 ml glass vials obtaining three analytical replicates. Nematodes were then pelleted by centrifugation and d17:1/12:0 sphingosylphosphoethanolamine (SPE; Sigma-Aldrich, 50 nM final concentration) added as internal standard. Lipids were extracted using LC-MS grade methanol, HPLC-grade chloroform (both Sigma-Aldrich) and Milli-Q water. After phase separation the lipid-containing chloroform phase was dried under nitrogen gas on a Techne Sample Concentrator. Lipids were re-suspended in 1 ml of acetonitrile: 10 mM ammonium acetate (pH 9.2) at a 95:5 (v:v) ratio. Samples were analyzed by liquid chromatography mass spectrometry (LC-MS) as follows: Five μl lipid extract were injected onto a Dionex UltiMate 3000RS UHPLC (Thermo Fisher Scientific) and separated on a hydrophilic interaction column (XBridge BEH amide XP column, Waters) according to their polar headgroup. The column was maintained at 30°C with a flow rate of 150 µl min^-1^. Samples were separated by a 15 min gradient from 95% (v:v) acetonitrile (Solvent A) to 30% (w:v) 10 mM ammonium acetate (pH 9.2, Solvent B) with 10 min equilibration between samples. Sample detection was carried out on an amaZon SL quadrupole ion trap MS (Bruker) in both positive and negative ion mode and fragmentation performed by the autoMS^n^ function in Compass HyStar (Bruker). We used the Bruker Compass software package for lipid data analysis: DataAnalysis for peak detection and lipid identification, and QuantAnalysis for quantification against the internal standard SPE. Peak integration was manually corrected where necessary. The abundance of each lipid was normalized against the internal standard SPE and expressed as relative abundance.

**Sample preparation and analysis for metabolomics.** Batches of 25 *Laxus oneistus* were extracted from the sand as described in Paredes et al., 2021. Sample collection to create triplicates lasted around 2 h, whereby the samples were always exposed to atmospheric oxygen. We aimed at comparing the metabolites present in the holobiont (*L. oneistus* and its ectosymbiont) and in the symbiont fraction (the dissociated bacteria). For the latter *Ca.* T. oneisti was dissociated from the nematodes by subjecting them to sonication for 1 min in 2 ml of seawater. The 2 ml nematode-free, ectosymbiont suspension was then centrifuged for 1 min at 14 000 x g to obtain *Ca.* T. oneisti pellets. Ectosymbiont and holobiont pellets were fixed in 500 µL of methanol and flash-frozen in liquid nitrogen and transported and stored at ‑80°C until further processing. Note that one biological replicate of the holobiont fraction was lost during transportation (Table S1).

For metabolite detection and quantification a gas-chromatography- mass spectrometry method was used. Nematode tissue or symbiont cells where separately extracted with an acetonitrile: methanol: water mixture as described previously (Liebeke and Bundy,2012) including a mechanical disruption of cells with ceramic beads in a bead beater (2x 30 sec, 4 m*s, Precellys, Bertin Instruments). The extracts where dried and later derivatized for GC-MS analysis (Liebeke and Puskás, 2019). The GC-MS analysis and metabolite identification was performed as described in Koch et al 2020). An in-house metabolite database of pure chemical standards was used for the identification and qualification of selected compounds.

**Data availability**. 'The mass spectrometry proteomics data have been deposited to the ProteomeXchange Consortium via the PRIDE (Perez-Riverol et al., 2019) partner repository (https://www.ebi.ac.uk/pride/) with the data set identifier PXD017709.
